# Quantification of metabolic niche occupancy dynamics in a Baltic Sea bacterial community

**DOI:** 10.1101/2022.08.08.502896

**Authors:** Jana C. Massing, Ashkaan Fahimipour, Carina Bunse, Jarone Pinhassi, Thilo Gross

## Abstract

Progress in molecular methods has enabled the monitoring of bacterial populations in time. Nevertheless, understanding community dynamics and its links with ecosystem functioning remains challenging due to the tremendous diversity of microorganisms. Conceptual frameworks that make sense of time-series of taxonomically-rich bacterial communities, regarding their potential ecological function, are needed. A key concept for organizing ecological functions is the niche, the set of strategies that enable a population to persist and define its impacts on the surroundings. Here we present a framework based on manifold learning, to organize genomic information into potentially occupied bacterial metabolic niches over time. We apply the method to re-construct the dynamics of putatively occupied metabolic niches using a long-term bacterial time-series from the Baltic Sea, the Linnaeus Microbial Observatory (LMO). The results reveal a relatively low-dimensional space of occupied metabolic niches comprising groups of taxa with similar functional capabilities. Time patterns of occupied niches were strongly driven by seasonality. Some metabolic niches were dominated by one bacterial taxon whereas others were occupied by multiple taxa, and this depended on season. These results illustrate the power of manifold learning approaches to advance our understanding of the links between community composition and functioning in microbial systems.

**Importance:** The increase in data availability of bacterial communities highlights the need for conceptual frameworks to advance our understanding of these complex and diverse communities alongside the production of such data. To understand the dynamics of these tremendously diverse communities, we need tools to identify overarching strategies and describe their role and function in the ecosystem in a comprehensive way. Here, we show that a manifold learning approach can coarse grain bacterial communities in terms of their metabolic strategies and that we can thereby quantitatively organize genomic information in terms of potentially occupied niches over time. This approach therefore advances our understanding of how fluctuations in bacterial abundances and species composition can relate to ecosystem functions and it can facilitate the analysis, monitoring and future predictions of the development of microbial communities.

## 1 Introduction

More than 60 years ago, Hutchinson formulated the paradox of plankton, expressing astonishment at the enormous diversity of organisms in the face of an apparently limited number of resources (1). Today, the estimated diversity of microbial species in the ocean extends this apparent contradiction to marine bacterial communities (2; 3). Over 40,000 marine microbial species have been detected so far (4; 5), and these microorganisms are critical for life in the oceans and on land because of their capacity to perform sophisticated and diverse chemical reactions, to drive major biogeochemical cycles (6), and to exhibit diverse interactions among each other and with macroorganisms (e.g. 7; 8). In addition to their huge diversity overall, marine bacterial communities undergo complex dynamic fluctuations in abundances and species compositions on the daily, monthly and annual scale (9; 10). In surface waters in temperate and polar regions, bacterial communities show strong seasonal patterns driven by changes in multiple interacting environmental features (9).

Due to the recent advances in modern molecular methods it has become possible to monitor bacterial community composition over time and establish short and long-term time series (examples in 11). These revealed for instance stability in average community composition in spite of strong variation on shorter scales (10), environmental selection as an important driver of seasonal community succession (12) and the significance of biological interactions among bacteria themselves, and between bacteria and other organisms e.g. phytoplankton, in shorter scale community dynamics (13).

Still, the tremendous diversity of bacteria poses a challenge for data analysis. Suppose for example that we sample a bacterial community and record the relative population densities for each detected taxon *e.g*., amplicon sequence variants, operational taxonomic units, species). In this case, the number of variables per sample is identical to the number of taxa. Mathematically we can say that the dimensionality of the data space equals the number of taxa. Dealing with such high-dimensional data is inherently difficult (the so-called curse of dimensionality) (14). In particular in high-dimensional spaces, comparisons between all but the most similar data points become so noisy that they hurt rather than help the analysis (15). Hence analysis can benefit from a coarse-graining step in which the dimensionality of the data set is reduced to a smaller number of variables. If done well this reduction yields new informative variables and also greatly reduces the noise in the data (16; 17).

When dealing with high-dimensional data spaces only a tiny portion of space is typically inhabited by data points. These data may approximately trace a curve, a curved surface, or some other comparatively low-dimensional object within the data space. Such objects in data-space are called data manifolds, and the task of locating them is known as manifold learning. Because the dimensionality of the manifold is lower than that of the embedding data space, manifold learning allows us to reduce the complexity of the data without losing information (18).

A widely used de-facto manifold learning method is principal component analysis (PCA) (19). PCA is a linear method that can only approximate the curved manifolds in the data by a flat surface. Therefore, it often works well in identifying global contrasts and for linearly distributed data. However, it cannot handle complex nonlinear data adequately. An alternative is offered by diffusion maps (20; 21; 22). The diffusion map identifies pairs of data points that are similar enough to be trusted even in the high dimensional space. Comparisons between any data points can then be made safely by measuring their distance on the network of all trusted comparisons. Mathematically the diffusion map is of similar complexity as PCA. However, unlike PCA it can identify curved manifolds and is immune from the curse of dimensionality. Like PCA, diffusion maps identify a new set of variables that describe the data in a coordinate system that follows the manifolds. Recent papers demonstrate that the application of diffusion maps to ecological data (22; 23; 24) yields new variables that can be interpreted as composite functional strategies. The diffusion map thus relates to the fundamental ecological niche space in a system (22; 25).

Here we apply diffusion maps to re-construct the metabolic niche space of a bacterial community from a long-term time-series in the Baltic Sea. We use the newly predicted variables to convert the taxonomic time-series into multiple strategy time-series and to quantify changes in functional diversity. Our results indicate that the diffusion map can reveal interpretable ecological strategies in the Baltic Sea bacterial community. This provides a quantitative framework to organize genomic information into potentially occupied ecological niches over time.

## 2 Results and Discussion

Our goal is to describe communities in terms of a coarse-grained functional coordinate system. Using diffusion maps, the coordinate axes that span this functional space can be found by a simple mathematical procedure (see Methods). The identified functional axes are new variables that are composites of functional capabilities of the analyzed bacterial community. These new variables are ordered according to their importance in structuring the dataset. Hence, in the following we refer to variable 1 as the most important, variable 2 the second most important and so on.

Moreover, the diffusion map assigns each species a score on these functional axes, i.e. a score for each new variable. Following Fahimipour and Gross (22) we interpret these new variables as effective composite metabolic strategies of the taxa. As each taxon from the dataset is assigned a score for each new variable, we can use these scores to order taxa from the most negative to the most positive entries for each variable. By analyzing the functional capabilities of exemplar taxa near the axis extrema, we can interpret the metabolic strategies that are described by the newly identified variables. For the interpretation, we divide each variable into positive and negative side because the taxa at the two sides can be characterized by different composite metabolic strategies.

Variable scores (from negative and positive side of the variable) can be used to convert the taxonomic time-series into multiple strategy time-series by calculating the abundance-weighted means of each variable side for each time-point (Figure 1). This enables us to observe the dynamics of community composition over time in terms of the dynamics of putatively occupied metabolic strategies, i.e. niches. Altogether the newly identified variables span the metabolic niche space of the analyzed community.

**Figure 1:**
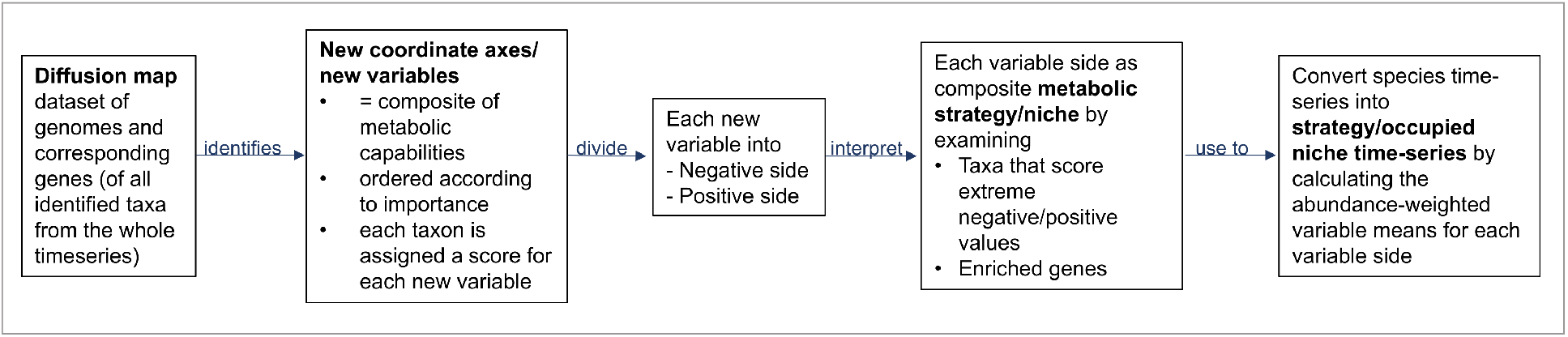
Overview of the procedure from diffusion mapping the dataset of genomes and genes to conversion of the species time-series into strategy time-series.

### 2.1 Important metabolic strategies

The most important variable identified via diffusion mapping, variable 1, separates primarily animal- or human-associated members of the Enterobacteriaceae *e.g*. close relatives of *Salmonella, Cronobacter* and *Yersinia* from all other taxa (Supplementary Table 2). A collection of 92 genomes from representatives of the Enterobacteriaceae score low values, whereas all other genomes are assigned values near zero (Figure 2 A). Such a variable in which the majority of species score close-to-zero is said to be a localized variable of the diffusion map (26), because it appears as a result of Anderson localization (27). In diffusion map results, the appearance of localized variables identifies a clear cluster of species that is well separated from the rest.

**Figure 2:**
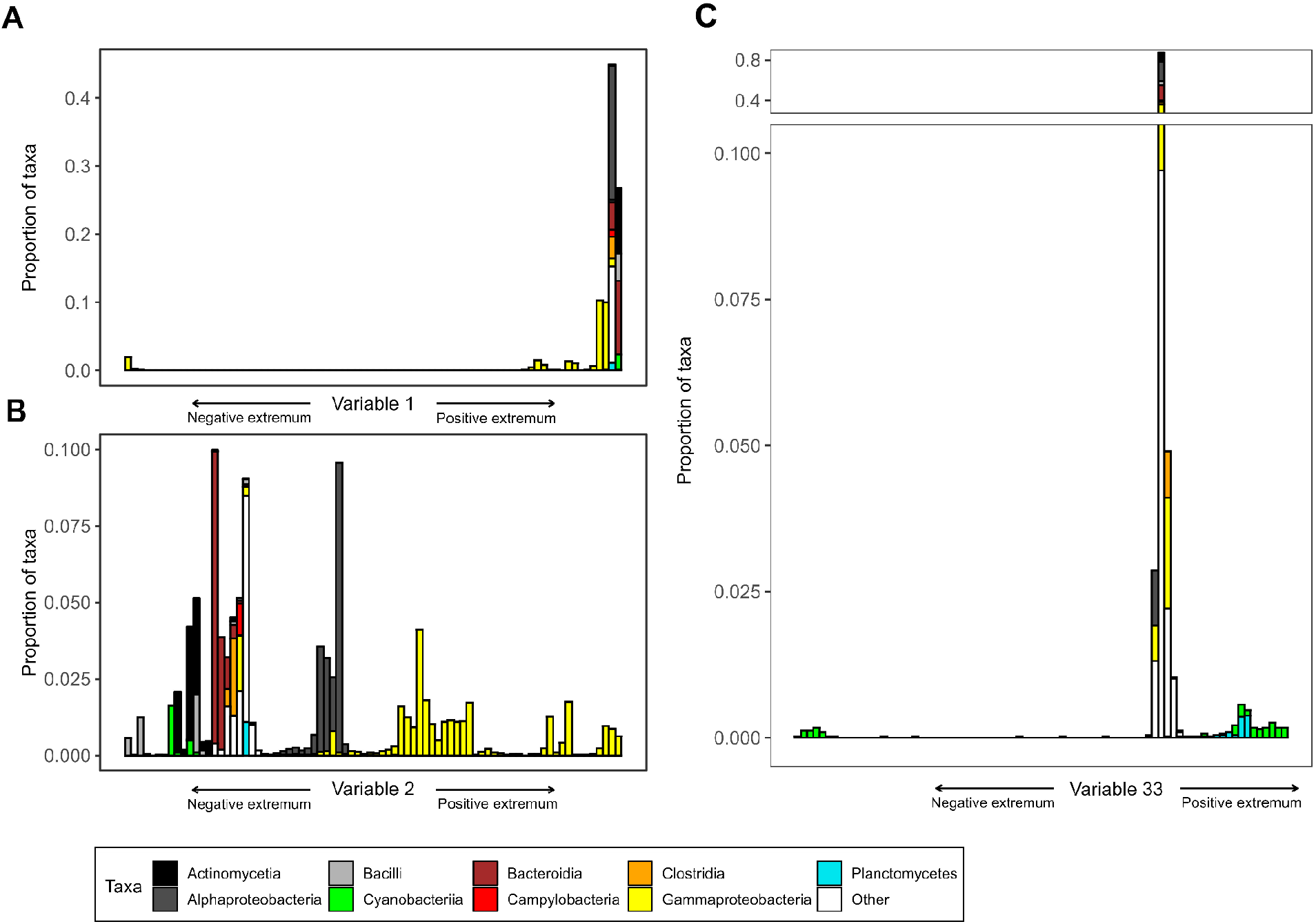
The ordering of taxa defined by variable 1 (A), variable 2 (B) and variable 33 (C) entries, from negative to positive (left to right). The taxonomic compositions corresponding to variable entries are shown for each of 80 equally spaced bins.

To reveal which metabolic capabilities characterize these taxa and distinguish them from all others, we identified genes that were over-represented in the genomes of the taxa scoring far-from-zero entries (see *Methods*). Genes encoding machinery for iron acquisition common in Enterobacteriaceae, such as the Enterobactin synthase component F (28; 29), genes responsible for the flagellar formation (30) and genes associated to biofilm formation (31) are enriched in these taxa (Supplementary Table 3).

Despite very low abundances of the respective taxa in the samples (mean relative abundances of 0.007 over all samples, see also SI material for abundance data), this variable appears first in the diffusion map. This may at first appear a surprising result given what is known about the ecological role of Enterobacteraceae in marine ecosystems. However, it makes sense that this group is so clearly separated from the other bacterial taxa due to well-known biases toward sequences of pathogenic taxa and genes involved in pathogenesis in global databases (e.g. 32). Consequently, these taxa are characterized as very different to other bacterial species in the marine community. Because the ‘full’ size fraction was sampled, we might have captured plankton pathogens (e.g. from zooplankton microbiomes or similar) that are very close relatives to these Enterobacteriaceae separated by the first diffusion variable. Hence, this variable separates the most different bacterial taxa in terms of their known gene composition from the rest of the community, demonstrating the power of the diffusion map method to reveal such differences and to identify biases in the dataset.

The diffusion map reveals further localized variables that represent relevant metabolic strategies for the Baltic Sea bacterial community, *e.g*. variable 4 negative, which separates the Cyanobacteria from all other taxa (Supplementary Figure 3). The cyanobacterial genomes are assigned large negative values, whereas all other taxa score positive or close-to-zero values. Genes that encode the subunits of photosystem I and photosystem II as well as associated cytochrome components and cyanobacterial-specific light-harvesting antennae (33) are among the enriched genes, indicating that this variable detects cyanobacterial photosynthesis (Supplementary Table 6). Supporting the findings of a previous study (22), the localized character of this variable reveals that cyanobacterial photosynthesis is a yes-or-no strategy, indicating that this photosynthetic lifestyle has wide-ranging metabolic consequences with distinct implications. For example, oxygenic photosynthetic lifestyle is expensive in terms of avoiding or repairing photoinhibition and -damage therefore many costly adaptations are necessary and the energy spent can not be invested into other metabolic pathways (34).

There are also variables that span a continuum of strategies, such as variable 2 and 3. In variable 2, marine host-associated Gammaproteobacteria, e.g. *Vibrio, Shewanella* and *Photobacterium*, are found at the positive extremum, whereas oligotrophic Gamma- and Alphaproteobacteria are assigned values close to zero (Figure 2 B). Among the most correlated capabilities for the taxa at the positive end of variable 2 are chemotaxis and response to various stressors (Supplementary Table 4). Variable 3 identifies the different strategies of marine Alphaproteobacteria: We find the Rhodobacteraceae and the Rhizobiales, known for their capability of utilizing a variety of carbon sources (35; 36), at the positive extremum (Supplementary Figure 2). Major enriched genes encode machinery for the utilization of various dissolved organic carbon compounds (Supplementary Table 5), *e.g*. phosphonate, acetate and urea that constitute exudates of phytoplankton (37). The streamlined genomes of the free-living Pelagibacterales and the obligate intracellular pathogens Rickettsiales score close-to-zero values. Hence we interpret variable 2 positive as metabolic strategy of marine host-associated Gammaproteobacteria and variable 3 positive as metabolic strategy of Alphaproteobacteria able of using a wide range of carbon sources.

Taxa that are grouped together in one variable can be separated by another variable. For example, Cyanobacteria group together in variable 4 negative, whereas they are split in variable 33: The Picocyanobacteria score extreme negative values, whereas the other cyanobacterial genomes group towards the positive side, with the heterocyst-forming family Nostocaceae scoring highest values (Figure 2 C and Supplementary Figure 6). The Enterobacterales that score highest in variable 2 are separated in variable 27, for which the family Shewanellaceae scores extreme negative values and the other enterobacterial families, *e.g*. the Vibrionaceae score positive values (Supplementary Figure 5). Variable 14 separates the Bacteroidota into anaerobic, intestinal Bacteroidota e.g. *Prevotella* (38) on the positive side and complex polysaccharide degraders e.g. *Flavobacterium* (39) on the negative side (Supplementary Figure 4). Enriched genes for the latter encode different CAZymes (40), responsible for the degradation of major plant cell wall components (Supplementary Table 7).

Diffusion mapping also identified strategies that group genera from different taxonomic groups together, we find for example bacteria that oxidize methyl groups and C1 compounds, such as methanol and formaldehyde from different families such as Beijerinckiaceae, Xanthobacteraceae, Acetobacteraceae at the negative end of variable 38 (Supplementary Figure 7). Most correlated genes (Supplementary Table 8) encode machinery for methanol and formaldehyde degradation (41). Another example is variable 43 positive that groups the non-spore forming sulfate-reducing bacteria from the families Desulfocapsaceae, Desulfobacteraceae, Desulfurivibrionaceae and others together (Supplementary Figure 8), enriched genes (Supplementary Table 9) are responsible for sulfate respiration (42).

### 2.2 Structure of the inferred niche space

The examples from above demonstrate that the diffusion map finds reasonable metabolic strategies of the bacterial community. Jointly these strategies define the metabolic niche space of the community and assign the taxa to specific coordinates in a multidimensional space. Using the PHATE procedure (15), we combined the diffusion variables to embed the data geometry in a low-dimensional visualization of the metabolic niche space (Figure 3). Although it is important to interpret low-dimensional embeddings cautiously, as in previous work, the strategy space of the Baltic Sea bacterial community appears as a tree-like structure comprising clusters of taxa featuring localized strategies and continuous branches (22) (Figure 3). The geometric structure implies large areas of the metabolic niche space that are unoccupied, reflecting either strategies that have yet to be discovered or strategies that are not feasible in the respective ecosystem or even in general (22).

**Figure 3:**
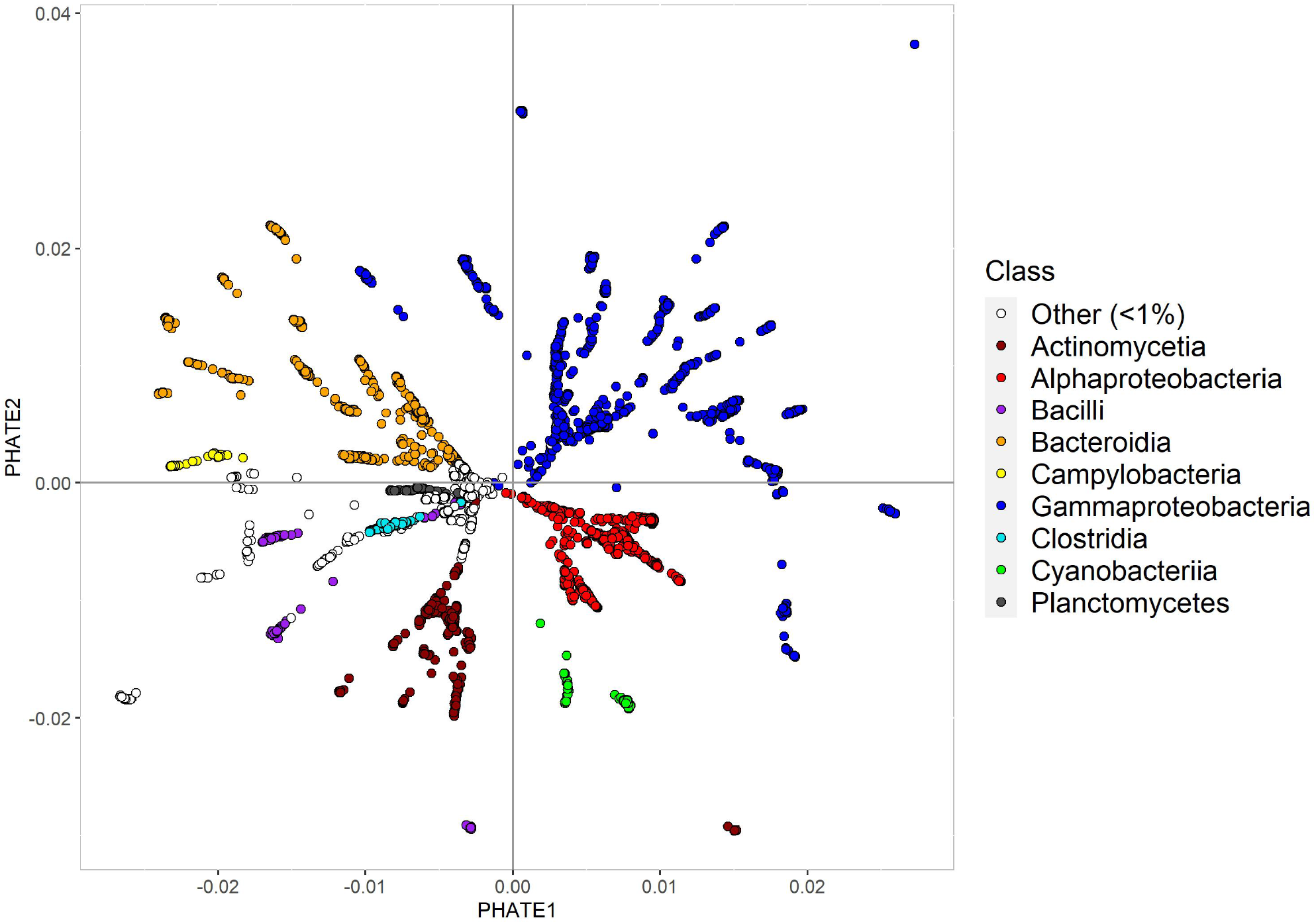
Two-dimensional embedding of diffusion variables created using the PHATE algorithm (15). Each point represents an individual genome that is coloured by taxonomic class.

The structure of the metabolic niche space roughly aligns with the phylogeny of the taxa. Uncultured taxa and the streamlined genomes of Patescibacter (43), Pelagibacterales (44) and Rickettsiales (45) form the core from which the branched structure emerges. The position of the streamlined genomes seems reasonable as these genomes retain mostly the key basic functions necessary for survival and reproduction that they share with many other organisms and they lack many of the more ‘specialized’ genes (46). The uncultured taxa group to the center either because they as well possess streamlined genomes or due to the lack of knowledge about their genes and respective functions. Therefore, their position in the metabolic niche space might change with further knowledge gained.

Taxa that are characterized by localized variables such as the Cyanobacteria are more distinct within the structure. These distinct clusters could indicate that complex, costly adaptions are necessary to follow the respective metabolic strategy and the machinery cannot easily be acquired for example through horizontal gene transfer (47). Localized strategies could also reflect that an intermediate strategy is not feasible, for instance due to certain trade-offs resulting from adopting the respective strategy (48).

Bacterial taxa associated with human diseases, like relatives of *Klebsiella, Mycobacterium, Staphylococcus* and *Fusobacterium*, group furthest away from the main structure and appear as clusters of dots in the periphery (Figure 3). The reason for these taxa to group away is probably the bias in global databases towards human pathogens and their functional capabilities (see above).

### 2.3 Metabolic strategies over time

Above, we saw that the new variables identified by diffusion mapping represent interpretable metabolic strategies of the sampled community. Taking the relative abundances of the taxa that map to different genomes into account, we calculated the abundance-weighted means of each diffusion variable side for each sampling time-point (see *Methods*). This enables us to observe how the occupation of metabolic niches or strategies change over time in the Baltic Sea bacterial community.

For example, the variable 4 negative, identified as cyanobacterial photosynthesis, reaches its highest abundance-weighted niche values in summer (Figure 4). Cyanobacteria are known to cause massive summer blooms in the Baltic Sea (49), supporting our interpretation that this strategy represents cyanobacterial photosynthesis. Separating the different cyanobacterial families reveals an early summer peak caused by the filamentous Nostocaceae and a plateau high of the niche values for the unicellular Cyanobiaceae increasing until the beginning of autumn (Figure 4). Utilization of nutrients from filamentous Cyanobacteria might fuel the metabolism of opportunistic picocyanobacteria (50).

**Figure 4:**
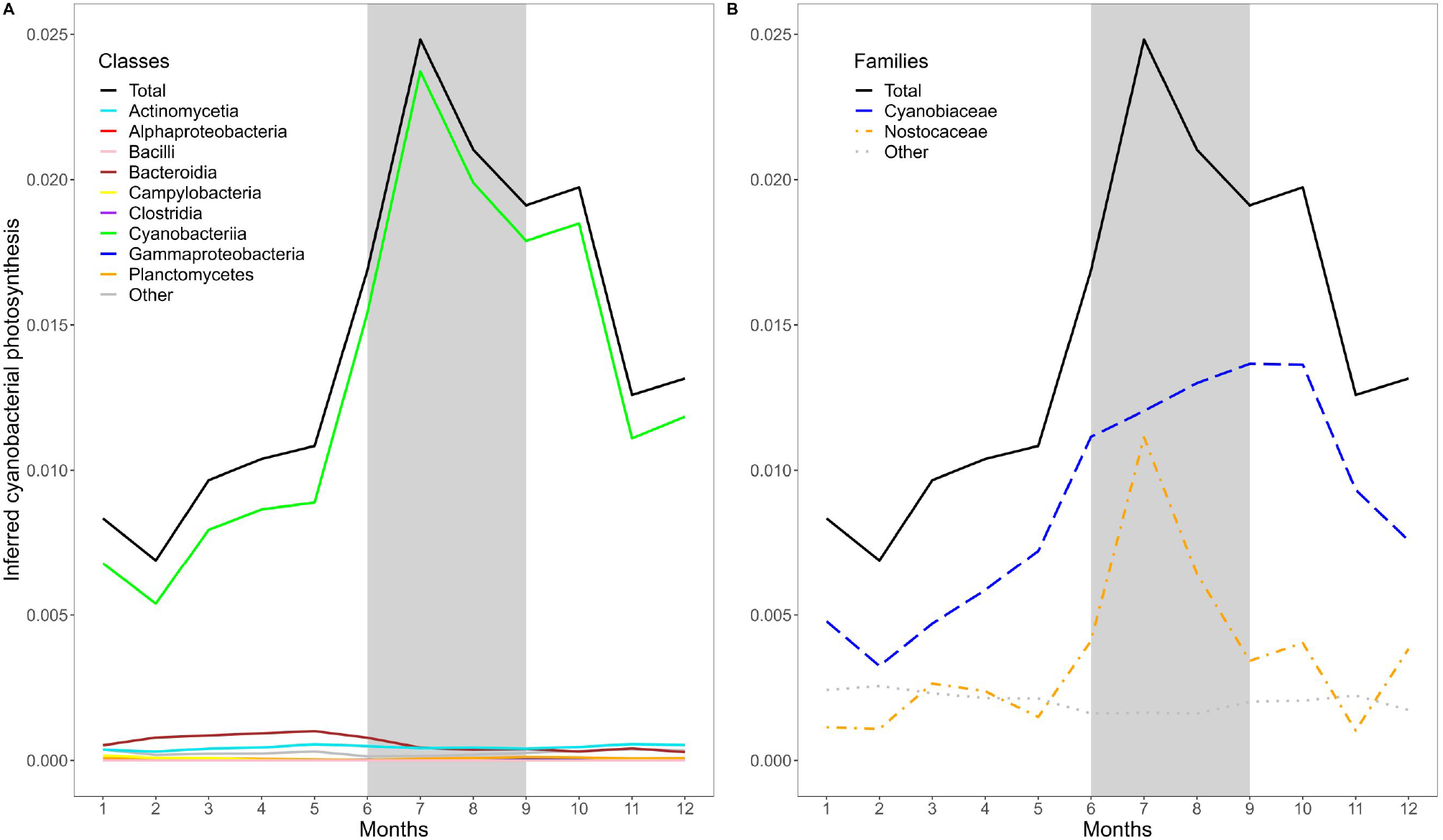
Abundance-weighted mean values of inferred cyanobacterial photosynthesis over the yearly cycle. Summer months are indicated by a gray background. Taxonomic class (A) and cyanobacterial taxonomic families (B) are color-coded.

Another strategy that is affected by seasonality is the catabolism of complex polysaccharides, identified in the variable 14 negative. Dominated by members of the Flavobacteriaceae, this strategy reaches highest abundance-weighted mean values in May (Figure 5 B), following the peak of the phytoplankton spring bloom (51). Large amounts of photosynthetic products, mainly polysaccharides, are exuded by phytoplankton (52) and Flavobacteria are adapted to use these high-molecular-weight molecules (39). In spring 2011 coinciding with the highest phytoplankton biomass (51), this strategy reaches highest values, compared to all other sampling years (Figure 5 A).

**Figure 5:**
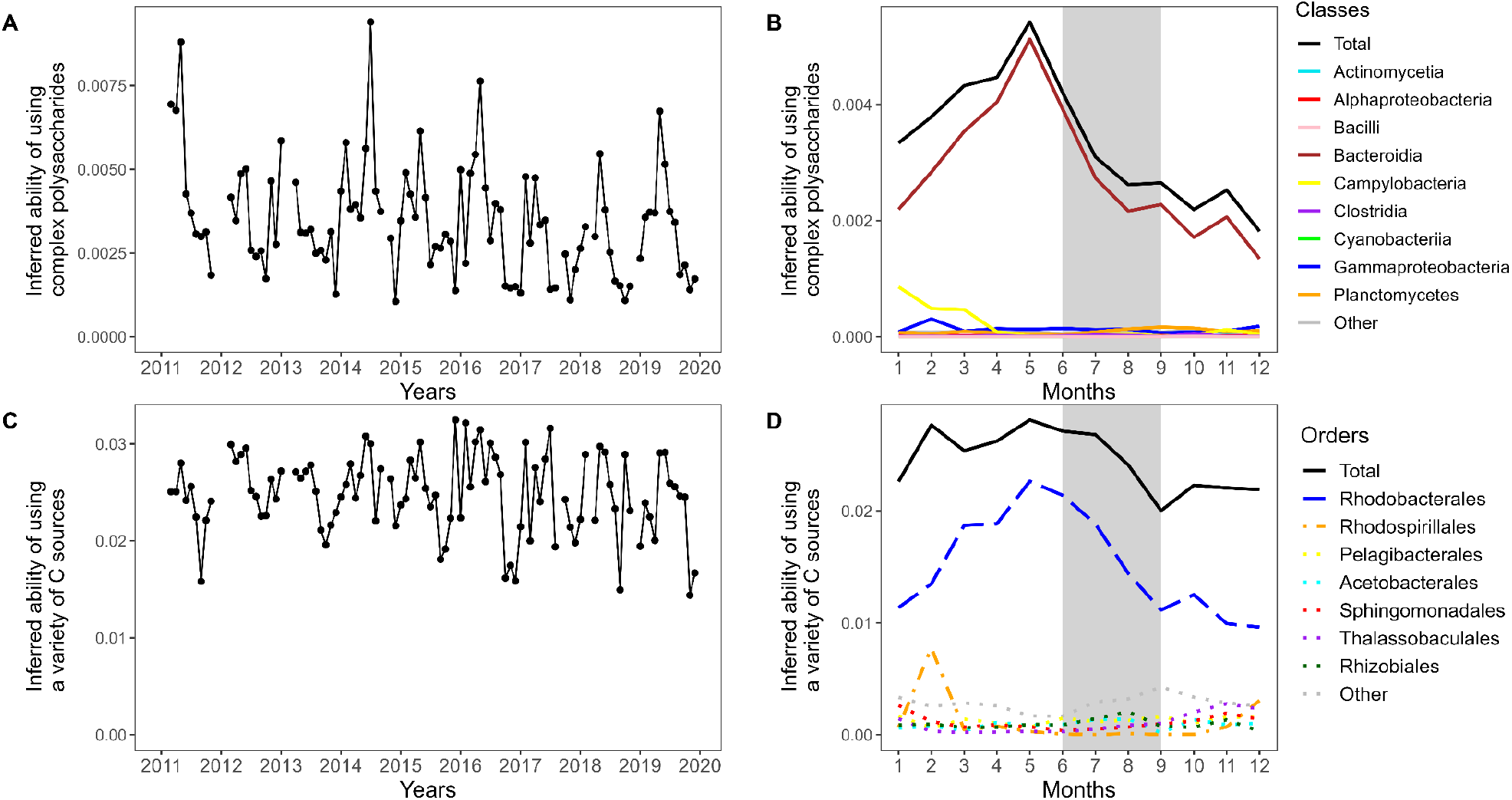
Abundance-weighted mean values of inferred ability of using complex polysaccharides (A, B) and inferred ability of using a variety of carbon sources(C,D) over the whole time period and over the yearly cycle. Summer months are indicated by a gray background. Taxonomic orders (B) and taxonomic classes (D) are color-coded.

Variable positive 3, describing the ability of utilizing a variety of carbon sources, also reaches highest values in May, but does not show a pronounced peak and instead decreases more slowly, reaching a minimum mean value in September (Figure 5 D). This strategy is dominated by the marine Rhodobacteraceae, that are crucial in processing low-molecular-weight phytoplankton-derived metabolites and characterized by their high trophic versatility (53; 54). In winter 2015/16 and 2016/17 elevated values of this variable positive 3 are driven by Rhodospirillales, especially of the genus *Thalassospira* (Figure 5 C). The latter is known for its ability to degrade polycylic aromatic hydrocarbons (PAHs) (55) and their appearance in the upper water column could be related to Major Baltic Inflow events (56) and subsequent winter mixing.

In contrast to the strategies we discussed above that were positively impacted by the seasonal phytoplankton spring bloom, variable negative 38 reaches its minimum in May, right after the phytoplankton bloom (Supplementary Figure 11). Describing the metabolic ability to oxidize methyl groups and C1 compounds, such as methanol and formaldehyde, this strategy is dominated by Alphaproteobacteria, especially *Pelagibacter* in winter and Planctomycetes in autumn in the Baltic Sea bacterial community. Marine dissolved organic carbon is a source for diverse C1 and methylated compounds, methanol constitutes a major fraction of oxygenated volatile organic chemicals and formaldehyde is omnipresent in seawater (41). The ability of using these compounds enables energy production from relatively abundant substrates in the water, but this ability is outcompeted when concentrations of phytoplankton-derived substrates increase in spring. Variable 43 positive describing non-spore forming sulfate reducers also reaches its minimum mean value in May (Supplementary Figure 12). Baltic Sea sulfate reducers of the phylum Desulfobacterota, inhabiting mostly sediments and oxygen-depleted waters (57; 58), drive the peak of this strategy in February, probably appearing in the upper water column due to strong winter mixing.

Comparing the strategy time-series to the environmental data obtained at LMO and the abundance-weighted mean values over the months, we can see a strong signal of seasonality (Figure 6): The variables on the left side of Figure 6 A are correlated to rather high nutrient concentrations and lower temperatures, hence winter conditions, whereas the variables on the right side of the heatmap are correlated to higher temperatures and chlorophyll a concentrations, hence summer conditions. Figure 6 B complements this picture: On the left side we find higher variable values in summer and autumn, while the variables on the right hand side show higher values in winter and spring.

**Figure 6:**
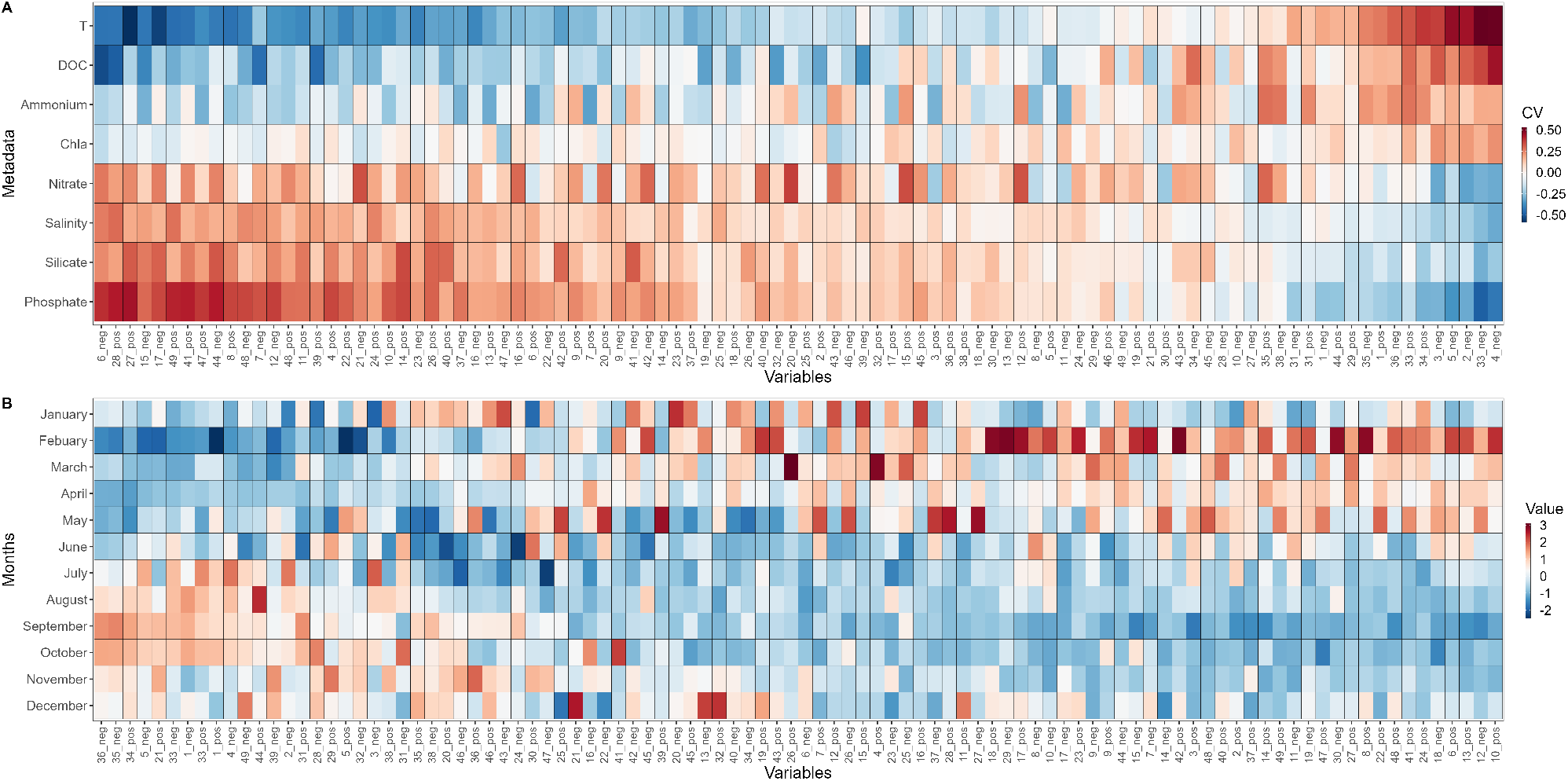
Heatmap of Spearman correlation coefficients (CV) between the first 49 variables, i.e. strategy time-series and the environmental variables (metadata) (A) and heatmap of abundance-weighted variable mean values of the first 49 variables for each month over the whole sampling period, standardized to mean = 0 and standard deviation = 1 for each variable side (B). T: temperature, DOC: dissolved organic carbon concentrations, Chla: chlorophyll a concentrations.

**Figure 7:**
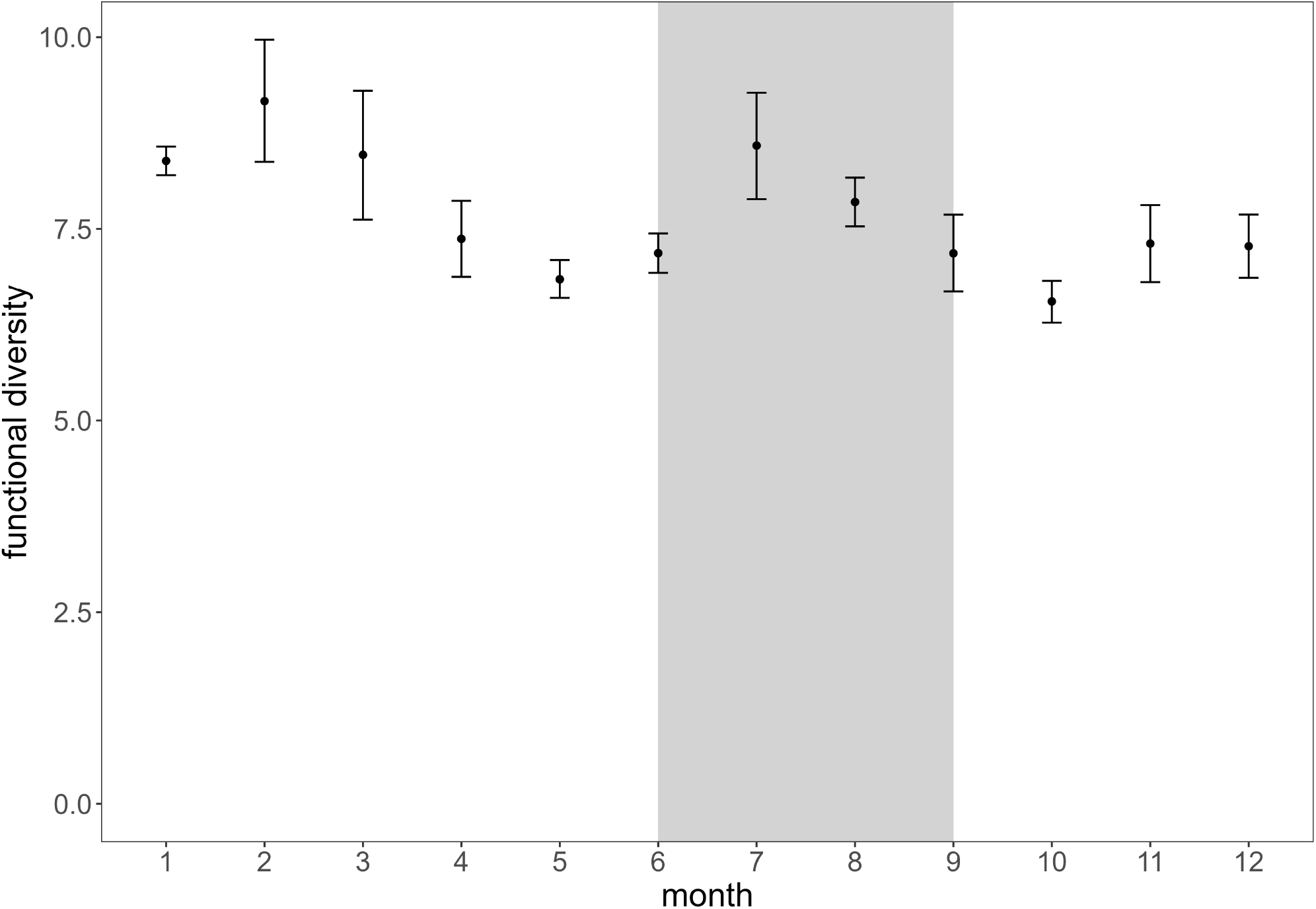
Mean and standard error of functional diversity estimation calculated as Rao index (23) for the sampled Baltic Sea bacterial community over the yearly cycle summarizing the years 2011-2019. Gray background indicates summer months.

### 2.4 Functional diversity

From the variables obtained via diffusion mapping we derived the diffusion distances between all pairs of species to quantify the functional diversity calculated as Rao index (23) (see Methods). The yearly cycle summarized over all sampling years shows highest functional diversity mean values in February and July, whereas it reaches lowest values in May and October (Fig. 7). This pattern is likely explained by the absence of the thermocline in winter, that causes deeper mixing of the water layers (59), resulting in bacterial communities from the former mesopelagic to be found also in surface waters (60; 61). Members of these communities can possess very different strategies due to their adaptation to deeper water layer or sediment conditions (61) and their taxonomic and functional diversity is in general higher than in surface communities (4), hence they increase the functional diversity when they are observed in the surface layers in winter. Functional diversity also benefits from increased diversity of nutrients available in winter, leading to an increased resource heterogeneity (62; 63). Lower values of functional diversity in May, following the phytoplankton bloom, could be explained by the strong dominance of strategies that relate to the utilization of phytoplankton-derived substrates during bloom phases as these compounds increase drastically in relative abundance (40; 39; 64).

## 3 Conclusion

In this paper we showed that relative abundance of prokaryotic populations obtained from amplicon sequencing data from monitoring datasets can be translated into potentially occupied ecological niches over time. The diffusion map of the bacterial capabilities detected a wide spectrum of interpretable metabolic strategies of the Baltic Sea bacterial community: From localized strategies such as cyanobacterial photosynthesis to continua of strategies such as degree of association with marine hosts and degree of trophic versatility, revealing both, strategies that align with phylogeny, strategies that differentiate closely-related taxa, and similar strategies among distantly-related taxa (22). The latter may reflect metabolic niche convergence (65) or horizontal gene transfer (e.g. 66).

Systematizing the genomic information via our diffusion map approach provided the possibility to express the changes in bacterial species abundances as quantitative changes in potential occupation of metabolic niches over time. These here called abundance-weighted strategy values showed a variety of patterns in time: Strategies displaying seasonal dynamics, e.g. increasing trends in summer or an increase following the phytoplankton bloom or higher values as a consequence of winter mixing as well as strategies showing interannual changes in time patterns. Impacts of single events, for example a very pronounced spring bloom or a Major Baltic Inflow event, were also reflected in the strategy time-series. Some functional strategies are clearly dominated by one bacterial group, whereas others are divided between bacterial groups often depending on season. Overall, seasonality seems to be a strong driver not only for phylogenetic bacterial composition (61; 60) but also for bacterial metabolic niche occupation and functional diversity of bacterial communities in the Baltic Sea. Seasonality is probably such an important driver also for metabolic strategies due to the interplay of seasonal changes in substrate availabilities together with changes in abiotic parameters that influence metabolic activities (67; 68; 60).

The first new variable identified by the diffusion map separates the metabolic strategies of pathogenic members of the Enterobacteriaceae from all other taxa. The reason for this variable showing up first is that the diffusion map probably detects the bias that bacterial pathogenic taxa and genes involved in pathogenesis are overrepresented in global databases (e.g. 32). This makes clear that a major limitation of this approach is our understanding of genes and their functions, implying also that the metabolic niche space might change due to further knowledge gained. For example, we might observe more branches representing currently undetected taxa or strategies and taxa currently grouped towards the center might shift further outwards and distances between taxa might change with increasing information. On the other hand this result also highlights the power of the diffusion map approach to objectively detect such biases in the data.

It should be noted that genes only encode the theoretical capabilities of a species (22), conceptually corresponding to the fundamental niche concept (25). It was however shown that functional genes can be used to predict the position of species along major niche gradients, outperforming the predictions based on phylogenetic information (69). Gowda et al. (70) also showed for the process of denitrification that it is possible to predict community metabolic dynamics from the presence and absence of genes in metagenomes. In comparison to their studies, our approach of assigning ASVs to species and obtaining their complete genomes from a database relies heavily on the quality of the databases and genome assemblages available. To overcome these shortcomings, the analysis of metagenomes is a desirable tool to improve our analysis by accounting also for within-species genome variability, the accessory genome and prophages that can play a role in shaping the strategies and traits of the respective organisms (71; 72; 73). Ideally, we would have all the genomes available from exactly the taxa from the habitat sampled, as we do not, we rely on a simple mapping scheme. In the future, deep shotgun metagenomic sequencing and long read technologies might be important for collecting the sorts of data that would make our method even more powerful.

We conclude that the diffusion map approach presented here enables us to coarse grain complex bacterial communities in terms of reasonable metabolic strategies and provides a quantitative framework to organize genomic information into potentially occupied metabolic niches over time. Thereby, this approach enables us to understand dynamics of community composition on different scales in terms of their impact on potentially occupied metabolic niches and to link genomic data to metabolic strategies enhancing our understanding of the relationships between community compositions and ecosystem functions.

## 4 Materials and Methods

### 4.1 Sampling data

Seawater samples were obtained from the Linnaeus Microbial Observatory (LMO) (N 56°55.854’, E 17°3.6420’) situated in the Western Baltic Proper approximately weekly during 2011-2013 and monthly during 2014-2019. Using 3 or 5 l Ruttner water samplers, water was sampled from 2 m depth at at ∼ 9 a.m. during each sampling occasion. Seawater was processed in the laboratory (Linnaeus University) where environmental parameters (temperature, salinity, chlorophyll a, DOC, nitrate and nitrite (together named as nitrate), phosphate, silicate and ammonium were analyzed as previously described (61; 51), Fridolfsson et al. in prep.). For microbial community composition, we filtered the seawater directly onto 0.22 µm SterivexTM cartridge filters (Millipore), or prefiltered onto 3 µm polycarbonate filters and subsequently on 0.22 µm SterivexTM cartridges (named 3-0.2 µm size fraction) using a persistaltic pump. We stored the filters in TE buffer at -80°C until DNA extraction using a phenol-chloroform method described by Boström et al. (74) and modified after Bunse et al. (75). We amplified the V3V4 region of the 16S rRNA gene using PCRs with the primer pair 341f-805r (76; 77). We analyzed the DNA concentrations with a Nanodrop or Qubit 2.0 Fluorometer (Life Technologies) and ran gel electrophoresis to confirm the amplicon specificity. Sample batches for sequencing were successively sent to the Science for Life Laboratory, Sweden on the Illumina MiSeq platform, resulting in 2 × 300 bp paired-end reads. For bioinformatic processing, we used the nf-core Ampliseq pipeline (78; 79) with the following software versions: nf-core/ampliseq = v1.2.0dev; Nextflow = v20.10.0; FastQC = v0.11.8; MultiQC = v1.9; Cutadapt = v2.8; QIIME2 = v2019.10.0. We used DADA2 (80) implemented in QIIME2 (81) and trimmed the sequences at forward 259 bp and reverse 199 bp before denoising. Of all LMO samples, we used all filter fractions for the niche space analysis but only the non-prefiltered 0.22 µm fraction for abundance estimates (see method description below). The LMO 16S rRNA raw data used in this study are deposited in the EMBL-EBI European Nucleotide Archive repository (https://www.ebi.ac.uk/ena) under accession numbers PRJEB52828, PRJEB52855, SRP048666, PRJEB52837, PRJEB40890, PRJEB52851, PRJEB52854, PRJEB52627, PRJEB52772, PRJEB52496, PRJEB52780, PRJEB52782, PRJEB52850.

### 4.2 Obtaining genomes and genes from ASV data

We conducted a BLAST sequence similarity search to match the ASV sequences obtained from 16S rRNA analysis from the time-series to the Genome Taxonomy Database (GTDB, https://data.gtdb.ecogenomic.org/, release 95 database). 21,102 (so 44%) of the 48,098 ASVs, could be matched well (similarity greater than 95%) to a genome from the GTDB. In terms of abundances, a mean of 82% (column sums) were good matches. From the well matched species we obtained the complete genomes from GTDB and NCBI (Refseq and GenBank). These 4,265 complete genomes were annotated using Prokka (82).

### 4.3 Diffusion mapping the strategy space

We diffusion mapped the genomes according to their similarity in known gene composition (absence-presence of genes) using the algorithm described in (16), applying the inverse of the hamming distance as a similarity measure. This method briefly consists of five steps. The starting point is the data matrix A with the dimensions *M* x *N*, where *M* =4,265 is the number of genomes and *N* = 15,361 is the number of annotated genes. First, the data is standardized so that each column has a mean of zero and a standard deviation of 1. Each row of our standardized data matrix can be interpreted as a vector of coordinates of the respective genome in a *N* -dimensional space. Second, we define similarities of two genomes as the inverse of the hamming distance between these vectors. The computed similarities form a *M*x*M* matrix, in which we set the diagonal elements to zero. We interpret the results as the weight matrix of the network with high values indicating close similarities between the respective genomes in terms of known gene composition. Third, we threshold the weight matrix, keeping only the top-25 highest similarities for each genome. From this weighted adjacency matrix we compute the corresponding row-normalized Laplacian matrix (step 4). In a final step, the eigenvectors and corresponding eigenvalues of this Laplacian are computed, defining new variables describing important variation in the dataset. The diffusion map identifies new variables spanning the strategy space of the analyzed bacteria and archaea and orders these variables according to their importance. Every genome is assigned a coordinate entry for each variable, so that we can order the corresponding taxa according to their entry, from most negative to most positive entries. Strategies of taxa that are assigned extreme entries along a variable can help interpret the meaning of each variable.

### 4.4 Translating ASV time-series into strategy time-series

For analysis, we separated the negative and positive values for each diffusion variable. To convert the species time-series into a strategy time-series, we calculated the weighted means of each diffusion variable for each sampling time-point, multiplying the eigenvector entries of each taxa by the normalized abundance of this taxa at this time-point.

### 4.5 Identifying over-represented genes

To identify the genes that were over-represented in the genomes of the taxa that themselves were assigned extreme entries along diffusion map variables, we used a permutational variant of the gene set enrichment analysis, GSEA (83). Genomes were ranked by the orderings specified by each diffusion variable. Gene set enrichment analysis was performed using the fgsea library in R (84; 85) with a Benjamini-Hochberg-adjusted (86) *P* value *<*0.01 used as a threshold for retaining genes corresponding to taxa that scored extreme values in the respective diffusion variable.

### 4.6 Estimating functional diversity

Functional diversity was estimated using the procedure described by Ryabov et al. (23). Briefly, the method uses the euclidean distances between species in the strategy space, rescaling the eigenvector elements according to their respective eigenvalue to quantify pairwise functional dissimilarities between all species. These newly defined diffusion distances can then be used to obtain the functional diversity of each sample, calculated as the Rao index.

### 4.7 Data processing

Data was processed with R 4.1.2 (84) and Julia 1.7.1 (87). The two dimensional mapping of the diffusion variables was made using the package phateR (15). Also the R packages ggplot2 (88), tidyverse (89), plyr (90), ggbreak (91), ggpubr (92), seriation (93) and reshape (94) were used.

## Supplemental Material file list

### Supplementary Tables

S1 Genomes that map to the 100 most abundant ASVs.

S2 Species that map to the 100 ASVs scoring most negative values in Variable 1.

S3 Top 100 over-represented annotated genes in the genomes of the taxa that receive the most negative entries on variable 1.

S4 Top 100 over-represented annotated genes in the genomes of the taxa that receive the most positive entries on variable 2.

S5 Top 100 over-represented annotated genes in the genomes of the taxa that receive the most positive entries on variable 3.

S6 Top 100 over-represented annotated genes in the genomes of the taxa that receive the most negative entries on variable 4.

S7 Top 100 over-represented annotated genes in the genomes of the taxa that receive the most negative entries on variable 14.

S8 Top 100 over-represented annotated genes in the genomes of the taxa that receive the most negative entries on variable 38.

S9 Top 100 over-represented annotated genes in the genomes of the taxa that receive the most positive entries on variable 43.

### Supplementary Figures

S1 Relative mean abundances of classes that map to the genomes obtained from amplicon sequencing data over the whole sampling period.

S2 The ordering of taxa defined by variable 3 entries.

S3 The ordering of taxa defined by variable 4 entries.

S4 The ordering of taxa defined by variable 14 entries.

S5 The ordering of taxa defined by variable 27 entries.

S6 The ordering of taxa defined by variable 33 entries.

S7 The ordering of taxa defined by variable 38 entries.

S8 The ordering of taxa defined by variable 43 entries.

S9 Abundance-weighted mean values of inferred ability of utilizing a variety of carbon sources over the yearly cycle.

S10 Abundance-weighted mean values of inferred ability of degrading complex polysaccharides over the yearly cycle.

S11 Abundance-weighted mean values of inferred ability of oxidizing methyl groups and C1 compounds over the yearly cycle.

S12 Abundance-weighted mean values of trait dominated by non-spore forming sulfate reducers over the yearly cycle.

## Acknowledgments

We thank the many people involved in LMO sampling and lab work over the years, especially Sabina Arnautovic, Anders Månsson, Emil Fridolfsson, Benjamin Pontiller and Kristofer Bergström. Furthermore, we acknowledge the help and assistance from Northern Offshore Services (NOS), M/V Provider crew, E.ON and RWE on the samplings. We acknowledge Daniel Lundin for bioinformatic support. The LMO research was supported by the Swedish Research Council FORMAS Strong Research environment EcoChange (Ecosystem dynamics in the Baltic Sea in a changing climate) to JP. CB, JM and TG were supported by the Helmholtz Institute for Functional Marine Biodiversity (HIFMB), a collaboration between the Alfred-Wegener-Institute, Helmholtz-Center for Polar and Marine Research, and the Carl-von-Ossietzky University Oldenburg, initially funded by the Ministry of Science and Culture of Lower Saxony (HIFMB Project) and the Volkswagen Foundation through the “Niedersächsisches Vorab” grant program (grant number ZN3285).We acknowledge support by the Open Access Publication Funds of Alfred-Wegener-Institut Helmholtz-Zentrum für Polar-und Meeresforschung.

## References

[1] G Evelyn Hutchinson. The paradox of the plankton. The American Naturalist, 95(882):137–145, 1961.

[2] Kenneth J Locey and Jay T Lennon. Scaling laws predict global microbial diversity. Proceedings of the National Academy of Sciences, 113(21):5970–5975, 2016.

[3] Stilianos Louca, Florent Mazel, Michael Doebeli, and Laura Wegener Parfrey. A census-based estimate of earth’s bacterial and archaeal diversity. PLoS biology, 17(2):e3000106, 2019.

[4] Shinichi Sunagawa, Luis Pedro Coelho, Samuel Chaffron, Jens Roat Kultima, Karine Labadie, Guillem Salazar, Bardya Djahanschiri, Georg Zeller, Daniel R Mende, Adriana Alberti, et al. Structure and function of the global ocean microbiome. Science, 348(6237):1261359, 2015.

[5] Weipeng Zhang, Wei Ding, Yong-Xin Li, Chunkit Tam, Salim Bougouffa, Ruojun Wang, Bite Pei, Hoyin Chiang, Pokman Leung, Yanhong Lu, et al. Marine biofilms constitute a bank of hidden microbial diversity and functional potential. Nature communications, 10(1):1–10, 2019.

[6] Paul G Falkowski, Tom Fenchel, and Edward F Delong. The microbial engines that drive earth’s biogeochemical cycles. science, 320(5879):1034–1039, 2008.

[7] Michael G Hadfield. Biofilms and marine invertebrate larvae: what bacteria produce that larvae use to choose settlement sites. Annual review of marine science, 3:453–470, 2011.

[8] David G Bourne, Kathleen M Morrow, and Nicole S Webster. Insights into the coral microbiome: underpinning the health and resilience of reef ecosystems. Annual Review of Microbiology, 70:317–340, 2016.

[9] Carina Bunse and Jarone Pinhassi. Marine bacterioplankton seasonal succession dynamics. Trends in microbiology, 25(6):494–505, 2017.

[10] Jed A Fuhrman, Jacob A Cram, and David M Needham. Marine microbial community dynamics and their ecological interpretation. Nature Reviews Microbiology, 13(3):133–146, 2015.

[11] Pier Luigi Buttigieg, Eduard Fadeev, Christina Bienhold, Laura Hehemann, Pierre Offre, and Antje Boetius. Marine microbes in 4d—using time series observation to assess the dynamics of the ocean microbiome and its links to ocean health. Current opinion in microbiology, 43:169–185, 2018.

[12] Adrià Auladell, Albert Barberán, Ramiro Logares, Esther Garcés, Josep M Gasol, and Isabel Ferrera. Seasonal niche differentiation among closely related marine bacteria. The ISME Journal, pages 1–12, 2021.

[13] David M Needham and Jed A Fuhrman. Pronounced daily succession of phytoplankton, archaea and bacteria following a spring bloom. Nature microbiology, 1(4):1–7, 2016.

[14] Richard Bellman. Dynamic programming. Princeton, USA: Princeton University Press, 1(2):3, 1957.

[15] Kevin R Moon, David van Dijk, Zheng Wang, Scott Gigante, Daniel B Burkhardt, William S Chen, Kristina Yim, Antonia van den Elzen, Matthew J Hirn, Ronald R Coifman, et al. Visualizing structure and transitions in high-dimensional biological data. Nature biotechnology, 37(12):1482–1492, 2019.

[16] Edmund Barter and Thilo Gross. Manifold cities: social variables of urban areas in the uk. Proceedings of the Royal Society A, 475(2221):20180615, 2019.

[17] Amin Ghafourian, Orestis Georgiou, Edmund Barter, and Thilo Gross. Wireless localization with diffusion maps. Scientific Reports, 10(1):1–10, 2020.

[18] Yunqian Ma and Yun Fu. Manifold learning theory and applications, volume 434. CRC press Boca Raton, 2012.

[19] Ian T Jolliffe and Jorge Cadima. Principal component analysis: a review and recent developments. Philosophical Transactions of the Royal Society A: Mathematical, Physical and Engineering Sciences, 374 (2065):20150202, 2016.

[20] Ronald R Coifman, Stephane Lafon, Ann B Lee, Mauro Maggioni, Boaz Nadler, Frederick Warner, and Steven W Zucker. Geometric diffusions as a tool for harmonic analysis and structure definition of data: Diffusion maps. Proceedings of the national academy of sciences, 102(21):7426–7431, 2005.

[21] Ronald R Coifman and Stéphane Lafon. Diffusion maps. Applied and computational harmonic analysis, 21 (1):5–30, 2006.

[22] Ashkaan K Fahimipour and Thilo Gross. Mapping the bacterial metabolic niche space. Nature communications, 11(1):4887, 2020.

[23] Alexey Ryabov, Bernd Blasius, Helmut Hillebrand, Irina Olenina, and Thilo Gross. Trait reconstruction and estimation of functional diversity from ecolocical monitoring data. arXiv preprint arXiv:2107.13792, 2021.

[24] Taylor Reiter, Rachel Montpetit, Ron Runnebaum, C Titus Brown, and Ben Montpetit. Charting shifts in saccharomyces cerevisiae gene expression across asynchronous time trajectories with diffusion maps. Mbio, 12(5):e02345–21, 2021.

[25] G Evelyn Hutchinson. Cold spring harbor symposium on quantitative biology. Concluding remarks, 22: 415–427, 1957.

[26] Amy Nyberg, Thilo Gross, and Kevin E Bassler. Mesoscopic structures and the laplacian spectra of random geometric graphs. Journal of Complex Networks, 3(4):543–551, 2015.

[27] Philip W Anderson. Absence of diffusion in certain random lattices. Physical review, 109(5):1492, 1958.

[28] J Ri Pollack and JB Neilands. Enterobactin, an iron transport compound from salmonella typhimurium. Biochemical and biophysical research communications, 38(5):989–992, 1970.

[29] IG O’brien and F Gibson. The structure of enterochelin and related 2, 3-dihydroxy-n-benzoyne conjugates from eschericha coli. Biochimica et Biophysica Acta (BBA)-General Subjects, 215(2):393–402, 1970.

[30] Phillip Aldridge and Kelly T Hughes. Regulation of flagellar assembly. Current opinion in microbiology, 5 (2):160–165, 2002.

[31] Joanna Domka, Jintae Lee, and Thomas K Wood. Ylih (bssr) and ycep (bsss) regulate escherichia coli k-12 biofilm formation by influencing cell signaling. Applied and environmental microbiology, 72(4):2449–2459, 2006.

[32] Grace A Blackwell, Martin Hunt, Kerri M Malone, Leandro Lima, Gal Horesh, Blaise TF Alako, Nicholas R Thomson, and Zamin Iqbal. Exploring bacterial diversity via a curated and searchable snapshot of archived dna sequences. PLoS biology, 19(11):e3001421, 2021.

[33] Patrick C Hallenbeck. Modern topics in the phototrophic prokaryotes: Metabolism, bioenergetics, and omics. Springer, 2017.

[34] John A Raven. The cost of photoinhibition. Physiologia Plantarum, 142(1):87–104, 2011.

[35] Haiwei Luo and Mary Ann Moran. How do divergent ecological strategies emerge among marine bacterioplankton lineages? Trends in microbiology, 23(9):577–584, 2015.

[36] Lucia Maria Carrareto Alves, Jackson Antônio Marcondes de Souza, Alessandro de Mello Varani, and EG de Macedo Lemos. The family rhizobiaceae. The Prokaryotes, pages 419–437, 2014.

[37] Karsten Zecher, Kristiane Rebecca Hayes, and Bodo Philipp. Evidence of interdomain ammonium crossfeeding from methylamine-and glycine betaine-degrading rhodobacteraceae to diatoms as a widespread interaction in the marine phycosphere. Frontiers in microbiology, 11, 2020.

[38] Adrian Tett, Edoardo Pasolli, Giulia Masetti, Danilo Ercolini, and Nicola Segata. Prevotella diversity, niches and interactions with the human host. Nature Reviews Microbiology, 19(9):585–599, 2021.

[39] Alison Buchan, Gary R LeCleir, Christopher A Gulvik, and José M González. Master recyclers: features and functions of bacteria associated with phytoplankton blooms. Nature Reviews Microbiology, 12(10): 686–698, 2014.

[40] Hanno Teeling, Bernhard M Fuchs, Dörte Becher, Christine Klockow, Antje Gardebrecht, Christin M Bennke, Mariette Kassabgy, Sixing Huang, Alexander J Mann, Jost Waldmann, et al. Substrate-controlled succession of marine bacterioplankton populations induced by a phytoplankton bloom. Science, 336(6081): 608–611, 2012.

[41] Jing Sun, Laura Steindler, J Cameron Thrash, Kimberly H Halsey, Daniel P Smith, Amy E Carter, Zachary C Landry, and Stephen J Giovannoni. One carbon metabolism in sar11 pelagic marine bacteria. PloS one, 6 (8):e23973, 2011.

[42] Sofia S Venceslau, Rita R Lino, and Ines AC Pereira. The qrc membrane complex, related to the alternative complex iii, is a menaquinone reductase involved in sulfate respiration. Journal of Biological Chemistry, 285 (30):22774–22783, 2010.

[43] Renmao Tian, Daliang Ning, Zhili He, Ping Zhang, Sarah J Spencer, Shuhong Gao, Weiling Shi, Linwei Wu, Ya Zhang, Yunfeng Yang, et al. Small and mighty: adaptation of superphylum patescibacteria to groundwater environment drives their genome simplicity. Microbiome, 8(1):1–15, 2020.

[44] Jana Grote, J Cameron Thrash, Megan J Huggett, Zachary C Landry, Paul Carini, Stephen J Giovannoni, and Michael S Rappé. Streamlining and core genome conservation among highly divergent members of the sar11 clade. MBio, 3(5):e00252–12, 2012.

[45] Alistair C Darby, Nam-Huyk Cho, Hans-Henrik Fuxelius, Joakim Westberg, and Siv GE Andersson. Intracellular pathogens go extreme: genome evolution in the rickettsiales. TRENDS in Genetics, 23(10): 511–520, 2007.

[46] Stephen J Giovannoni, J Cameron Thrash, and Ben Temperton. Implications of streamlining theory for microbial ecology. The ISME journal, 8(8):1553–1565, 2014.

[47] Robert E Blankenship. Early evolution of photosynthesis. Plant physiology, 154(2):434–438, 2010.

[48] Thomas Ferenci. Trade-off mechanisms shaping the diversity of bacteria. Trends in microbiology, 24(3): 209–223, 2016.

[49] Markus V Lindh and Jarone Pinhassi. Sensitivity of bacterioplankton to environmental disturbance: a review of baltic sea field studies and experiments. Frontiers in Marine Science, 5:361, 2018.

[50] Mireia Bertos-Fortis, Hanna M Farnelid, Markus V Lindh, Michele Casini, Agneta Andersson, Jarone Pinhassi, and Catherine Legrand. Unscrambling cyanobacteria community dynamics related to environmental factors. Frontiers in microbiology, 7:625, 2016.

[51] Carina Bunse, Stina Israelsson, Federico Baltar, Mireia Bertos-Fortis, Emil Fridolfsson, Catherine Legrand, Elin Lindehoff, Markus V Lindh, Sandra Martínez-García, and Jarone Pinhassi. High frequency multi-year variability in baltic sea microbial plankton stocks and activities. Frontiers in microbiology, 9:3296, 2019.

[52] Marco Mühlenbruch, Hans-Peter Grossart, Falk Eigemann, and Maren Voss. Mini-review: Phytoplankton-derived polysaccharides in the marine environment and their interactions with heterotrophic bacteria. Environmental microbiology, 20(8):2671–2685, 2018.

[53] Frank Xavier Ferrer-González, Brittany Widner, Nicole R Holderman, John Glushka, Arthur S Edison, Elizabeth B Kujawinski, and Mary Ann Moran. Resource partitioning of phytoplankton metabolites that support bacterial heterotrophy. The ISME journal, 15(3):762–773, 2021.

[54] Ryan J Newton, Laura E Griffin, Kathy M Bowles, Christof Meile, Scott Gifford, Carrie E Givens, Erinn C Howard, Eric King, Clinton A Oakley, Chris R Reisch, et al. Genome characteristics of a generalist marine bacterial lineage. The ISME journal, 4(6):784–798, 2010.

[55] Taishi Tsubouchi, Yukari Ohta, Takuma Haga, Keiko Usui, Yasuhiro Shimane, Kozue Mori, Akiko Tanizaki, Akiko Adachi, Kiwa Kobayashi, Kiyotaka Yukawa, et al. Thalassospira alkalitolerans sp. nov. and thalassospira mesophila sp. nov., isolated from a decaying bamboo sunken in the marine environment, and emended description of the genus thalassospira. International journal of systematic and evolutionary microbiology, 64 (Pt 1):107–115, 2014.

[56] Benjamin Bergen, Michael Naumann, Daniel PR Herlemann, Ulf Gräwe, Matthias Labrenz, and Klaus Jürgens. Impact of a major inflow event on the composition and distribution of bacterioplankton communities in the baltic sea. Frontiers in Marine Science, page 383, 2018.

[57] Adrien Vetterli, Kirsi Hyytiäinen, Minttu Ahjos, Petri Auvinen, Lars Paulin, Susanna Hietanen, and Elina Leskinen. Seasonal patterns of bacterial communities in the coastal brackish sediments of the gulf of finland, baltic sea. Estuarine, Coastal and Shelf Science, 165:86–96, 2015.

[58] VA Korneeva, NV Pimenov, AV Krek, TP Tourova, and AL Bryukhanov. Sulfate-reducing bacterial communities in the water column of the gdansk deep (baltic sea). Microbiology, 84(2):268–277, 2015.

[59] Jan Hinrich Reissmann, Hans Burchard, Rainer Feistel, Eberhard Hagen, Hans Ulrich Lass, Volker Mohrholz, Günther Nausch, Lars Umlauf, and Gunda Wieczorek. Vertical mixing in the baltic sea and consequences for eutrophication–a review. Progress in Oceanography, 82(1):47–80, 2009.

[60] Daniel PR Herlemann, Daniel Lundin, Anders F Andersson, Matthias Labrenz, and Klaus Jürgens. Phylogenetic signals of salinity and season in bacterial community composition across the salinity gradient of the baltic sea. Frontiers in Microbiology, 7:1883, 2016.

[61] Markus V Lindh, Johanna Sjöstedt, Anders F Andersson, Federico Baltar, Luisa W Hugerth, Daniel Lundin, Saraladevi Muthusamy, Catherine Legrand, and Jarone Pinhassi. Disentangling seasonal bacterioplankton population dynamics by high-frequency sampling. Environmental microbiology, 17(7):2459–2476, 2015.

[62] Angelika Rieck, Daniel PR Herlemann, Klaus Jürgens, and Hans-Peter Grossart. Particle-associated differ from free-living bacteria in surface waters of the baltic sea. Frontiers in Microbiology, 6:1297, 2015.

[63] Eric Raes, Jennifer Tolman, Dhwani Desai, Jenni-Marie Ratten, Jackie Zorz, Brent M Robicheau, Diana Haider, and Julie LaRoche. Seasonal bacterial niche structures and chemolithoautotrophic ecotypes in a north atlantic fjord. doi:doi.org/10.21203/rs.3.rs-1469763/v1, 2022.

[64] Jakob Pernthaler. Competition and niche separation of pelagic bacteria in freshwater habitats. Environmental microbiology, 19(6):2133–2150, 2017.

[65] Eric R Pianka, Laurie J Vitt, Nicolás Pelegrin, Daniel B Fitzgerald, and Kirk O Winemiller. Toward a periodic table of niches, or exploring the lizard niche hypervolume. The American Naturalist, 190(5): 601–616, 2017.

[66] J Peter Gogarten and Jeffrey P Townsend. Horizontal gene transfer, genome innovation and evolution. Nature Reviews Microbiology, 3(9):679–687, 2005.

[67] Christopher S Ward, Cheuk-Man Yung, Katherine M Davis, Sara K Blinebry, Tiffany C Williams, Zackary I Johnson, and Dana E Hunt. Annual community patterns are driven by seasonal switching between closely related marine bacteria. The ISME journal, 11(6):1412–1422, 2017.

[68] Michael Seidel, Marcus Manecki, Daniel PR Herlemann, Barbara Deutsch, Detlef Schulz-Bull, Klaus Jürgens, and Thorsten Dittmar. Composition and transformation of dissolved organic matter in the baltic sea. Frontiers in Earth Science, 5:31, 2017.

[69] Johannes Alneberg, Christin Bennke, Sara Beier, Carina Bunse, Christopher Quince, Karolina Ininbergs, Lasse Riemann, Martin Ekman, Klaus Jürgens, Matthias Labrenz, et al. Ecosystem-wide metagenomic binning enables prediction of ecological niches from genomes. Communications biology, 3(1):1–10, 2020.

[70] Karna Gowda, Derek Ping, Madhav Mani, and Seppe Kuehn. Genomic structure predicts metabolite dynamics in microbial communities. Cell, 2022.

[71] Natalia García-García, Javier Tamames, Alexandra M Linz, Carlos Pedrós-Alió, and Fernando Puente-Sánchez. Microdiversity ensures the maintenance of functional microbial communities under changing environmental conditions. The ISME journal, 13(12):2969–2983, 2019.

[72] Ana Zhu, Shinichi Sunagawa, Daniel R Mende, and Peer Bork. Inter-individual differences in the gene content of human gut bacterial species. Genome biology, 16(1):1–13, 2015.

[73] Kathryn Forcone, Felipe H Coutinho, Giselle S Cavalcanti, and Cynthia B Silveira. Prophage genomics and ecology in the family rhodobacteraceae. Microorganisms, 9(6):1115, 2021.

[74] Kjärstin H Boström, Karin Simu, Åke Hagström, and Lasse Riemann. Optimization of dna extraction for quantitative marine bacterioplankton community analysis. Limnology and Oceanography: Methods, 2(11): 365–373, 2004.

[75] Carina Bunse, Mireia Bertos-Fortis, Ingrid Sassenhagen, Sirje Sildever, Conny Sjöqvist, Anna Godhe, Susanna Gross, Anke Kremp, Inga Lips, Nina Lundholm, et al. Spatio-temporal interdependence of bacteria and phytoplankton during a baltic sea spring bloom. Frontiers in microbiology, 7:517, 2016.

[76] Daniel PR Herlemann, Matthias Labrenz, Klaus Jürgens, Stefan Bertilsson, Joanna J Waniek, and Anders F Andersson. Transitions in bacterial communities along the 2000 km salinity gradient of the baltic sea. The ISME journal, 5(10):1571–1579, 2011.

[77] Luisa W Hugerth, Hugo A Wefer, Sverker Lundin, Hedvig E Jakobsson, Mathilda Lindberg, Sandra Rodin, Lars Engstrand, and Anders F Andersson. Degeprime, a program for degenerate primer design for broad-taxonomic-range pcr in microbial ecology studies. Applied and environmental microbiology, 80(16): 5116–5123, 2014.

[78] Philip A Ewels, Alexander Peltzer, Sven Fillinger, Harshil Patel, Johannes Alneberg, Andreas Wilm, Maxime Ulysse Garcia, Paolo Di Tommaso, and Sven Nahnsen. The nf-core framework for community-curated bioinformatics pipelines. Nature biotechnology, 38(3):276–278, 2020.

[79] Daniel Straub, Nia Blackwell, Adrian Langarica-Fuentes, Alexander Peltzer, Sven Nahnsen, and Sara Kleindienst. Interpretations of environmental microbial community studies are biased by the selected 16s rrna (gene) amplicon sequencing pipeline. Frontiers in Microbiology, 11:550420, 2020.

[80] Benjamin J Callahan, Paul J McMurdie, Michael J Rosen, Andrew W Han, Amy Jo A Johnson, and Susan P Holmes. Dada2: High-resolution sample inference from illumina amplicon data. Nature methods, 13(7): 581–583, 2016.

[81] Evan Bolyen, Jai Ram Rideout, Matthew R Dillon, Nicholas A Bokulich, Christian C Abnet, Gabriel A Al-Ghalith, Harriet Alexander, Eric J Alm, Manimozhiyan Arumugam, Francesco Asnicar, et al. Reproducible, interactive, scalable and extensible microbiome data science using qiime 2. Nature biotechnology, 37(8): 852–857, 2019.

[82] Torsten Seemann. Prokka: rapid prokaryotic genome annotation. Bioinformatics, 30(14):2068–2069, 2014.

[83] Aravind Subramanian, Pablo Tamayo, Vamsi K Mootha, Sayan Mukherjee, Benjamin L Ebert, Michael A Gillette, Amanda Paulovich, Scott L Pomeroy, Todd R Golub, Eric S Lander, et al. Gene set enrichment analysis: a knowledge-based approach for interpreting genome-wide expression profiles. Proceedings of the National Academy of Sciences, 102(43):15545–15550, 2005.

[84] R Core Team. R: A Language and Environment for Statistical Computing. R Foundation for Statistical Computing, Vienna, Austria, 2020. URL https://www.R-project.org/.

[85] Gennady Korotkevich, Vladimir Sukhov, and Alexey Sergushichev. Fast gene set enrichment analysis. bioRxiv, 2019. doi: 10.1101/060012. URL http://biorxiv.org/content/early/2016/06/20/060012.

[86] Yoav Benjamini and Yosef Hochberg. Controlling the false discovery rate: a practical and powerful approach to multiple testing. Journal of the Royal statistical society: series B (Methodological), 57(1):289–300, 1995.

[87] Jeff Bezanson, Alan Edelman, Stefan Karpinski, and Viral B Shah. Julia: A fresh approach to numerical computing. SIAM Review, 59(1):65–98, 2017. doi: 10.1137/141000671. URL https://epubs.siam.org/doi/10.1137/141000671.

[88] Hadley Wickham. ggplot2: Elegant Graphics for Data Analysis. Springer-Verlag New York, 2016. ISBN 978-3-319-24277-4. URL https://ggplot2.tidyverse.org.

[89] Hadley Wickham, Mara Averick, Jennifer Bryan, Winston Chang, Lucy D’Agostino McGowan, Romain François, Garrett Grolemund, Alex Hayes, Lionel Henry, Jim Hester, Max Kuhn, Thomas Lin Pedersen, Evan Miller, Stephan Milton Bache, Kirill Müller, Jeroen Ooms, David Robinson, Dana Paige Seidel, Vitalie Spinu, Kohske Takahashi, Davis Vaughan, Claus Wilke, Kara Woo, and Hiroaki Yutani. Welcome to the tidyverse. Journal of Open Source Software, 4(43):1686, 2019. doi: 10.21105/joss.01686.

[90] Hadley Wickham. The split-apply-combine strategy for data analysis. Journal of Statistical Software, 40(1): 1–29, 2011. URL https://www.jstatsoft.org/v40/i01/.

[91] Shuangbin Xu, Meijun Chen, Tingze Feng, Li Zhan, Lang Zhou, and Guangchuang Yu. Use ggbreak to effectively utilize plotting space to deal with large datasets and outliers. Frontiers in Genetics, 12:774846, 2021. doi: 10.3389/fgene.2021.774846.

[92] Alboukadel Kassambara. ggpubr: ‘ggplot2’ Based Publication Ready Plots, 2020. URL https://rpkgs.datanovia.com/ggpubr/. R package version 0.4.0.999.

[93] Michael Hahsler, Kurt Hornik, and Christian Buchta. Getting things in order: An introduction to the r package seriation. Journal of Statistical Software, 25(3):1–34, March 2008. ISSN 1548-7660. doi: 10.18637/jss.v025.i03.

[94] Hadley Wickham. Reshaping data with the reshape package. Journal of Statistical Software, 21(12), 2007. URL http://www.jstatsoft.org/v21/i12/paper.

