## Supplementary Material for "Quantification of metabolic niche occupancy dynamics in a Baltic Sea bacterial community"

#### Supplementary Tables

**Table S1:** Genomes that map to the 100 most abundant ASVs obtained from amplicon sequencing data in terms of relative mean abundance over the whole sampling period. Genome, taxonomic information, mean and maximum abundance over the whole sampling period are provided.

| Genome | Class | Family | Species | Mean abundance | Max abundance |
|---|---|---|---|---|---|
| GCF_003011885.1 | Cyanobacteriia | Cyanobiaceae | Cyanobium_A usitatum | 0.1150 | 0.7222 |
| GCF_002252665.1 | Cyanobacteriia | Cyanobiaceae | Cyanobium_A sp002252665 | 0.0533 | 0.2779 |
| GCA_003569125.1 | Acidimicrobiia | Ilumatobacteraceae | BACL27 sp003569125 | 0.0488 | 0.3656 |
| GCA_001593825.1 | Cyanobacteriia | Nostocaceae | Aphanizomenon_B flosaquae | 0.0400 | 0.5437 |
| GCF_000173115.1 | Bacteroidia | Flavobacteriaceae | MAG-120531 sp000173115 | 0.0397 | 0.5443 |
| GCA_002358295.1 | Gammaproteobacteria | D2472 | D2472 sp002358345 | 0.0287 | 0.2754 |
| GCA_003569145.1 | Actinomycetia | Nanopelagicaceae | MAG-120802 sp003569145 | 0.0286 | 0.2757 |
| GCA_001438235.1 | Alphaproteobacteria | Rhodobacteraceae | UBA10365 sp003536295 | 0.0235 | 0.1846 |
| GCA_002405515.1 | Planctomycetes | UBA1268 | UBA4655 sp002405515 | 0.0226 | 0.1789 |
| GCA_007280255.1 | Planctomycetes | UBA1268 | QWOQ01 sp003669585 | 0.0213 | 0.2010 |
| GCA_001437765.1 | Acidimicrobiia | Ilumatobacteraceae | UBA3006 sp002367695 | 0.0208 | 0.2445 |
| GCA_002325485.1 | Bacteroidia | Flavobacteriaceae | BACL21 sp002694465 | 0.0201 | 0.3039 |
| GCA_002340585.1 | Gammaproteobacteria | Porticoccaceae | HTCC2207 sp001438605 | 0.0191 | 0.2454 |
| GCF_001485105.1 | Actinomycetia | Streptomycetaceae | Streptomyces acidiscabies | 0.0163 | 0.3385 |
| GCA_002711735.1 | Acidimicrobiia | Ilumatobacteraceae | Ilumatobacter_A sp002711735 | 0.0142 | 0.2166 |
| GCF_000496475.1 | Gammaproteobacteria | Burkholderiaceae | RS62 sp000496475 | 0.0139 | 0.1293 |
| GCF_000257665.1 | Actinomycetia | Microbacteriaceae | Aquiluna sp000257665 | 0.0129 | 0.2494 |
| GCF_000312705.1 | Cyanobacteriia | Nostocaceae | LE011-02 sp000312705 | 0.0122 | 0.5288 |
| GCF_002287885.2 | Actinomycetia | Nanopelagicaceae | Nanopelagicus limnes | 0.0114 | 0.0733 |
| GCF_000242915.1 | Campylobacteria | Sulfurimonadaceae | Sulfurimonas gotlandica | 0.0107 | 0.6154 |
| GCA_002746305.1 | Bacteroidia | UBA9320 | UBA9320 sp002746305 | 0.0100 | 0.0928 |
| GCA_002430225.1 | Actinomycetia | Microbacteriaceae | Pontimonas sp001438965 | 0.0093 | 0.2095 |
| GCF_002252705.1 | Cyanobacteriia | Cyanobiaceae | Vulcanococcus limneticus | 0.0092 | 0.1075 |
| GCA_000750175.1 | Alphaproteobacteria | Pelagibacteraceae | IMCC9063 sp000750175 | 0.0090 | 0.1093 |
| GCA_002340845.1 | Gammaproteobacteria | Methylophilaceae | BACL14 sp002384685 | 0.0079 | 0.0679 |
| GCA_001438645.1 | Gammaproteobacteria | Methylophilaceae | BACL14 sp002384685 | 0.0076 | 0.0700 |
| GCA_001438145.1 | Gammaproteobacteria | Pseudohongiellaceae | OM182 sp001438145 | 0.0074 | 0.0646 |
| GCF_002284895.1 | Actinomycetia | Nanopelagicaceae | Planktophila sp002284895 | 0.0074 | 0.0575 |
| GCA_000485495.1 | Actinomycetia | Nanopelagicaceae | AAA044-D11 sp000485495 | 0.0074 | 0.0642 |
| GCA_002170165.1 | Bacteroidia | BACL11 | TMED123 sp002170165 | 0.0072 | 0.0501 |
| GCF_900129545.1 | Bacteroidia | Flavobacteriaceae | Flavobacterium fluvii | 0.0060 | 0.2432 |
| GCF_001983935.1 | Planctomycetes | Planctomycetaceae | Fuerstia marisgermanicae | 0.0058 | 0.1676 |
| GCA_001438305.1 | Bacteroidia | Schleiferiaceae | TMED14 sp001438205 | 0.0056 | 0.0693 |
| GCF_002252635.1 | Cyanobacteriia | Cyanobiaceae | WH-5701 sp002252635 | 0.0055 | 0.0868 |
| GCA_004292795.1 | Bacteroidia | Microscillaceae | RDXI01 sp004292795 | 0.0054 | 0.1049 |
| GCF_006491595.1 | Bacteroidia | Flavobacteriaceae | Flavobacterium jejuense | 0.0052 | 0.0682 |
| GCF_002943715.1 | Bacteroidia | Flavobacteriaceae | Polaribacter filamentus | 0.0052 | 0.1530 |
| GCF_000299115.1 | Alphaproteobacteria | HIMB59 | HIMB59 sp000299115 | 0.0051 | 0.0616 |
| GCF_900114485.1 | Alphaproteobacteria | Rhodobacteraceae | Loktanella salsilacus | 0.0049 | 0.0452 |
| GCA_004379135.1 | Acidimicrobiia | Ilumatobacteraceae | Casp-actino8 sp004379135 | 0.0048 | 0.0220 |
| GCA_003249095.1 | Cyanobacteriia | Microcystaceae | Snowella sp003249095 | 0.0047 | 0.1042 |
| GCF_002631185.1 | Alphaproteobacteria | Acetobacteraceae | Roseomonas rhizosphaerae | 0.0047 | 0.3294 |
| GCA_000738435.1 | Alphaproteobacteria | Rhodobacteraceae | Planktomarina temperata | 0.0046 | 0.1321 |
| GCF_003096315.1 | Gammaproteobacteria | Burkholderiaceae | Achromobacter insuavis | 0.0046 | 0.4719 |
| GCA_003284275.1 | Alphaproteobacteria | Pelagibacteraceae | Pelagibacter_A sp003284275 | 0.0045 | 0.0547 |
| GCF_002252675.1 | Cyanobacteriia | Cyanobiaceae | Cyanobium sp002252675 | 0.0044 | 0.0434 |
| GCF_002954645.1 | Bacteroidia | Flavobacteriaceae | Polaribacter gangjinensis | 0.0043 | 0.0744 |
| GCF_002101315.1 | Alphaproteobacteria | Pelagibacteraceae | Pelagibacter sp002101315 | 0.0040 | 0.0398 |
| GCF_002115755.1 | Alphaproteobacteria | Thalassospiraceae | Thalassospira mesophila | 0.0039 | 0.2429 |

| | | | | | |
|---|---|---|---|---|---|
| GCA_002733565.1 | Gammaproteobacteria | Psychromonadaceae | Moritella sp000170855 | 0.0039 | 0.0718 |
| GCF_002288225.1 | Actinomycetia | Nanopelagicaceae | Planktophila dulcis | 0.0039 | 0.0286 |
| GCA_002428815.1 | Gammaproteobacteria | Porticoccaceae | HTCC2207 sp001438605 | 0.0038 | 0.0499 |
| GCF_000590925.1 | Alphaproteobacteria | Rhodobacteraceae | Roseicyclus elongatum | 0.0037 | 0.0555 |
| GCA_002346275.1 | Gammaproteobacteria | Halieaceae | IMCC3088 sp003520285 | 0.0037 | 0.0802 |
| GCF_002940745.1 | Bacteroidia | Flavobacteriaceae | Hanstruepera crassostreae | 0.0036 | 0.0623 |
| GCF_003335085.1 | Bacteroidia | Flavobacteriaceae | Polaribacter sp003335085 | 0.0036 | 0.1667 |
| GCF_000699505.1 | Actinomycetia | Microbacteriaceae | Rhodoluna lacicola | 0.0035 | 0.0265 |
| GCA_003149555.1 | Actinomycetia | Microbacteriaceae | Aquiluna sp003149555 | 0.0034 | 0.0827 |
| GCA_004379115.1 | Actinomycetia | S36-B12 | Mxb001 sp004379115 | 0.0034 | 0.0716 |
| GCA_001438005.1 | Verrucomicrobiae | UBA3015 | UBA3015 sp001438005 | 0.0032 | 0.0291 |
| GCA_002346225.1 | Bacteroidia | BACL12 | UBA11426 sp002346225 | 0.0031 | 0.1118 |
| GCF_003003055.1 | Gammaproteobacteria | Burkholderiaceae | SCGC-AAA027-K21 sp003003055 | 0.0031 | 0.0317 |
| GCF_002284855.1 | Actinomycetia | Nanopelagicaceae | Planktophila sp002284855 | 0.0029 | 0.0379 |
| GCA_002863125.1 | Bacteroidia | UA16 | UA16 sp002863125 | 0.0029 | 0.0250 |
| GCF_002284915.1 | Actinomycetia | Nanopelagicaceae | IMCC26077 sp002284915 | 0.0029 | 0.0279 |
| GCA_001438165.1 | Bacteroidia | Schleiferiaceae | TMED14 sp002381225 | 0.0029 | 0.0274 |
| GCF_001457835.1 | Clostridia | Ezakiellaceae | Fenollaria timonensis | 0.0029 | 0.1789 |
| GCF_000152785.1 | Alphaproteobacteria | Rhodobacteraceae | Yoonia vestfoldensis_A | 0.0028 | 0.0290 |
| GCA_002292365.1 | Bacteroidia | Cyclobacteriaceae | UBA4465 sp002292365 | 0.0028 | 0.0194 |
| GCF_001439695.1 | Gammaproteobacteria | Pseudomonadaceae | Pseudomonas_E veronii | 0.0028 | 0.1510 |
| GCA_000421325.1 | Alphaproteobacteria | AAA536-G10 | AAA536-G10 sp000421325 | 0.0027 | 0.0406 |
| GCA_003208775.1 | Cyanobacteriia | Cyanobiaceae | Synechococcus_C sp002500205 | 0.0026 | 0.1927 |
| GCA_003671255.1 | Planctomycetes | Gemmataceae | UBA969 sp003671255 | 0.0025 | 0.0379 |
| GCA_007093895.1 | Gammaproteobacteria | Enterobacteriaceae | Salmonella enterica | 0.0024 | 0.0168 |
| GCF_003011125.1 | Cyanobacteriia | Cyanobiaceae | Synechococcus_D lacustris | 0.0024 | 0.0380 |
| GCF_004337435.1 | Actinomycetia | Streptomycetaceae | Streptomyces sp004337435 | 0.0024 | 0.0120 |
| GCA_002167745.1 | Gammaproteobacteria | SG8-40 | UBA3031 sp002167745 | 0.0023 | 0.0320 |
| GCA_900618205.1 | Gammaproteobacteria | Burkholderiaceae | Bordetella trematum | 0.0023 | 0.1770 |
| GCF_000173095.1 | Bacteroidia | Flavobacteriaceae | MS024-2A sp000173095 | 0.0023 | 0.0428 |
| GCF_000797465.1 | Bacteroidia | Flavobacteriaceae | Psychroserpens jangbogonensis | 0.0021 | 0.0377 |
| GCA_002690755.1 | Phycisphaerae | SM1A02 | UBA12014 sp002690755 | 0.0021 | 0.0255 |
| GCA_002480055.1 | Gammaproteobacteria | Porticoccaceae | HTCC2207 sp002335945 | 0.0020 | 0.0215 |
| GCF_000143825.1 | Actinomycetia | Mycobacteriaceae | Corynebacterium genitalium_A | 0.0020 | 0.0926 |
| GCF_006385135.1 | Alphaproteobacteria | Emcibacteraceae | Emcibacter_A congregatus | 0.0020 | 0.0176 |
| GCF_002284875.1 | Actinomycetia | Nanopelagicaceae | Planktophila sp002284875 | 0.0019 | 0.0111 |
| GCA_002697205.1 | Gammaproteobacteria | HTCC2089 | GCA-2697205 sp002697205 | 0.0019 | 0.0172 |
| GCA_003045825.1 | Bacteroidia | Schleiferiaceae | UBA10364 sp003045825 | 0.0019 | 0.0476 |
| GCA_000762985.1 | Actinomycetia | Mycobacteriaceae | Mycobacterium rufum | 0.0018 | 0.0245 |
| GCA_002282055.1 | Bacteroidia | Sphingobacteriaceae | Daejeonella sp002257025 | 0.0018 | 0.0238 |
| GCA_002499015.1 | Poseidoniia | Poseidoniaceae | MGIIa-L1 sp002499015 | 0.0018 | 0.0959 |
| GCF_000176015.1 | Alphaproteobacteria | Rhodobacteraceae | Pseudorhodobacter_B sp000176015 | 0.0018 | 0.0476 |
| GCF_006937785.1 | Cyanobacteriia | Pseudanabaenaceae | Pseudanabaena sp006937785 | 0.0017 | 0.1092 |
| GCF_001623485.1 | Cyanobacteriia | Nostocaceae | Nodularia spumigena | 0.0016 | 0.0348 |
| GCF_000171835.1 | Alphaproteobacteria | Thalassobaculaceae | BAL199 sp000171835 | 0.0015 | 0.0138 |
| GCF_000156155.1 | Gammaproteobacteria | Methylophilaceae | BACL14 sp000156155 | 0.0015 | 0.0159 |
| GCA_002733945.1 | Campylobacteria | Sulfurimonadaceae | Sulfurimonas sp002733945 | 0.0015 | 0.0763 |
| GCF_003856375.1 | Bacteroidia | Crocinitomicaceae | Fluviicola sp003856375 | 0.0015 | 0.0449 |
| GCF_900110395.1 | Alphaproteobacteria | Reyranellaceae | Reyranella sp900110395 | 0.0015 | 0.0222 |
| GCF_002368115.1 | Cyanobacteriia | Nostocaceae | Dolichospermum_A compactum | 0.0014 | 0.0579 |
| GCF_900100865.1 | Actinomycetia | Microbacteriaceae | Aquiluna sp900100865 | 0.0014 | 0.0188 |

**Table S2:** Species that map to the 100 ASVs scoring most negative values in variable 1.

| **Species** | **Value** | **Genome** | **Class** | **Family** |
|---|---|---|---|---|
| Salmonella enterica | -0.09769 | GCA_007094035.1 | Gammaproteobacteria | Enterobacteriaceae |
| Salmonella enterica | -0.09769 | GCA_007093895.1 | Gammaproteobacteria | Enterobacteriaceae |
| Salmonella enterica | -0.09763 | GCA_007093765.1 | Gammaproteobacteria | Enterobacteriaceae |
| Salmonella enterica | -0.09756 | GCF_003548795.1 | Gammaproteobacteria | Enterobacteriaceae |
| Cronobacter sakazakii | -0.09754 | GCF_002094495.1 | Gammaproteobacteria | Enterobacteriaceae |
| Cronobacter sakazakii | -0.09754 | GCF_002094665.1 | Gammaproteobacteria | Enterobacteriaceae |
| Cronobacter sakazakii | -0.09754 | GCF_002977865.1 | Gammaproteobacteria | Enterobacteriaceae |
| Cronobacter turicensis | -0.09754 | GCF_002976545.1 | Gammaproteobacteria | Enterobacteriaceae |
| Cronobacter sakazakii | -0.09753 | GCA_002094675.1 | Gammaproteobacteria | Enterobacteriaceae |
| Cronobacter sakazakii | -0.09753 | GCF_002094645.1 | Gammaproteobacteria | Enterobacteriaceae |
| Cronobacter sakazakii | -0.09752 | GCF_002977315.1 | Gammaproteobacteria | Enterobacteriaceae |
| Cronobacter malonaticus | -0.09752 | GCF_002978245.1 | Gammaproteobacteria | Enterobacteriaceae |
| Cronobacter malonaticus | -0.09752 | GCF_002978235.1 | Gammaproteobacteria | Enterobacteriaceae |
| Cronobacter sakazakii | -0.09751 | GCF_002094575.1 | Gammaproteobacteria | Enterobacteriaceae |
| Cronobacter sakazakii | -0.09751 | GCF_002976775.1 | Gammaproteobacteria | Enterobacteriaceae |
| Cronobacter malonaticus | -0.09751 | GCF_002978375.1 | Gammaproteobacteria | Enterobacteriaceae |
| Cronobacter sakazakii | -0.09751 | GCF_002094585.1 | Gammaproteobacteria | Enterobacteriaceae |
| Cronobacter sakazakii | -0.09751 | GCA_002976965.2 | Gammaproteobacteria | Enterobacteriaceae |
| Cronobacter dublinensis | -0.09751 | GCF_002979155.1 | Gammaproteobacteria | Enterobacteriaceae |
| Cronobacter malonaticus | -0.09751 | GCF_002978545.1 | Gammaproteobacteria | Enterobacteriaceae |
| Cronobacter malonaticus | -0.09751 | GCF_002978185.1 | Gammaproteobacteria | Enterobacteriaceae |
| Cronobacter sakazakii | -0.09751 | GCF_002977405.1 | Gammaproteobacteria | Enterobacteriaceae |
| Cronobacter sakazakii | -0.09751 | GCF_002094475.1 | Gammaproteobacteria | Enterobacteriaceae |
| Cronobacter malonaticus | -0.0975 | GCF_002978535.1 | Gammaproteobacteria | Enterobacteriaceae |
| Cronobacter sakazakii | -0.0975 | GCF_002976735.1 | Gammaproteobacteria | Enterobacteriaceae |
| Cronobacter sakazakii | -0.0975 | GCF_002976795.1 | Gammaproteobacteria | Enterobacteriaceae |
| Cronobacter sakazakii | -0.0975 | GCA_002977005.2 | Gammaproteobacteria | Enterobacteriaceae |
| Escherichia coli | -0.09749 | GCA_002078275.1 | Gammaproteobacteria | Enterobacteriaceae |
| Cronobacter sakazakii | -0.09749 | GCF_002977155.1 | Gammaproteobacteria | Enterobacteriaceae |
| Escherichia coli | -0.09749 | GCF_001268585.1 | Gammaproteobacteria | Enterobacteriaceae |
| Escherichia coli | -0.09749 | GCF_001269185.1 | Gammaproteobacteria | Enterobacteriaceae |
| Escherichia coli | -0.09748 | GCF_004523105.1 | Gammaproteobacteria | Enterobacteriaceae |
| Escherichia coli | -0.09748 | GCF_005889645.1 | Gammaproteobacteria | Enterobacteriaceae |

| | | | | |
|---|---|---|---|---|
| Escherichia coli | -0.09748 | GCF_001268685.1 | Gammaproteobacteria | Enterobacteriaceae |
| Escherichia coli | -0.09748 | GCF_002007165.1 | Gammaproteobacteria | Enterobacteriaceae |
| Escherichia coli | -0.09748 | GCF_002959275.1 | Gammaproteobacteria | Enterobacteriaceae |
| Cronobacter sakazakii | -0.09748 | GCF_002978035.1 | Gammaproteobacteria | Enterobacteriaceae |
| Cronobacter sakazakii | -0.09747 | GCF_002978105.1 | Gammaproteobacteria | Enterobacteriaceae |
| Cronobacter dublinensis | -0.09747 | GCA_002978875.2 | Gammaproteobacteria | Enterobacteriaceae |
| Cronobacter dublinensis | -0.09747 | GCF_002978655.1 | Gammaproteobacteria | Enterobacteriaceae |
| Enterobacter sp. | -0.09746 | GCF_000534395.1 | Gammaproteobacteria | Enterobacteriaceae |
| Scandinavium goeteborgense | -0.09744 | GCF_004361715.1 | Gammaproteobacteria | Enterobacteriaceae |
| Scandinavium goeteborgense | -0.09743 | GCA_003935895.2 | Gammaproteobacteria | Enterobacteriaceae |
| Klebsiella pneumoniae | -0.09742 | GCF_003967395.1 | Gammaproteobacteria | Enterobacteriaceae |
| Cronobacter dublinensis | -0.09742 | GCF_002978855.1 | Gammaproteobacteria | Enterobacteriaceae |
| Klebsiella quasivariicola | -0.09742 | GCF_002269255.1 | Gammaproteobacteria | Enterobacteriaceae |
| Klebsiella variicola | -0.09742 | GCF_001033575.1 | Gammaproteobacteria | Enterobacteriaceae |
| Klebsiella quasipneumoniae | -0.09742 | GCF_002853635.1 | Gammaproteobacteria | Enterobacteriaceae |
| Klebsiella pneumoniae | -0.09742 | GCF_004127885.1 | Gammaproteobacteria | Enterobacteriaceae |
| Citrobacter freundii | -0.09741 | GCA_001686345.1 | Gammaproteobacteria | Enterobacteriaceae |
| Enterobacter cloacae | -0.09741 | GCF_001562175.1 | Gammaproteobacteria | Enterobacteriaceae |
| Enterobacter sp. | -0.0974 | GCF_000493015.1 | Gammaproteobacteria | Enterobacteriaceae |
| Cronobacter sakazakii | -0.0974 | GCF_002977115.1 | Gammaproteobacteria | Enterobacteriaceae |
| Citrobacter koseri | -0.09739 | GCF_002393245.1 | Gammaproteobacteria | Enterobacteriaceae |
| Klebsiella pneumoniae | -0.09739 | GCF_003227185.1 | Gammaproteobacteria | Enterobacteriaceae |
| Klebsiella pneumoniae | -0.09739 | GCF_004127515.1 | Gammaproteobacteria | Enterobacteriaceae |
| Pantoea sp. | -0.09737 | GCF_002920175.1 | Gammaproteobacteria | Enterobacteriaceae |
| Klebsiella grimontii | -0.09736 | GCA_902159485.1 | Gammaproteobacteria | Enterobacteriaceae |
| Enterobacter sp. | -0.09736 | GCA_007035975.1 | Gammaproteobacteria | Enterobacteriaceae |
| Pantoea sp. | -0.09736 | GCF_002313185.2 | Gammaproteobacteria | Enterobacteriaceae |
| Pantoea sp. | -0.09736 | GCF_003813865.1 | Gammaproteobacteria | Enterobacteriaceae |
| Buttiauxella izardii | -0.09735 | GCF_003601925.1 | Gammaproteobacteria | Enterobacteriaceae |
| Klebsiella grimontii | -0.09735 | GCA_902158675.1 | Gammaproteobacteria | Enterobacteriaceae |
| Lelliottia nimipressuralis | -0.09734 | GCF_004402045.1 | Gammaproteobacteria | Enterobacteriaceae |
| Pantoea stewartii | -0.09732 | GCF_001310295.1 | Gammaproteobacteria | Enterobacteriaceae |
| Pantoea sp. | -0.09729 | GCF_000963985.1 | Gammaproteobacteria | Enterobacteriaceae |
| Rahnella aquatilis | -0.09729 | GCF_000735505.1 | Gammaproteobacteria | Enterobacteriaceae |
| Buttiauxella sp. | -0.09728 | GCF_003675305.1 | Gammaproteobacteria | Enterobacteriaceae |
| Cronobacter dublinensis | -0.09727 | GCF_002978705.2 | Gammaproteobacteria | Enterobacteriaceae |
| Erwinia sp. | -0.09726 | GCF_002752575.1 | Gammaproteobacteria | Enterobacteriaceae |
| Rouxiella chamberiensis | -0.09724 | GCF_000951135.1 | Gammaproteobacteria | Enterobacteriaceae |
| Erwinia sp. | -0.09724 | GCF_004551645.1 | Gammaproteobacteria | Enterobacteriaceae |
| Enterobacter sp. | -0.09721 | GCF_000277545.1 | Gammaproteobacteria | Enterobacteriaceae |
| Gamma proteobacterium | -0.09715 | GCA_000335795.1 | Gammaproteobacteria | Enterobacteriaceae |
| Rahnella aquatilis | -0.09711 | GCA_003956145.2 | Gammaproteobacteria | Enterobacteriaceae |
| Rahnella woolbedingensis | -0.09709 | GCF_003602095.1 | Gammaproteobacteria | Enterobacteriaceae |
| Ewingella americana | -0.09702 | GCF_006438725.1 | Gammaproteobacteria | Enterobacteriaceae |
| Ewingella americana | -0.09696 | GCF_900451015.1 | Gammaproteobacteria | Enterobacteriaceae |
| Serratia quinivorans | -0.09689 | GCF_900457075.1 | Gammaproteobacteria | Enterobacteriaceae |
| Atlantibacter hermannii | -0.09685 | GCF_900635495.1 | Gammaproteobacteria | Enterobacteriaceae |
| Serratia proteamaculans | -0.09685 | GCF_004684015.1 | Gammaproteobacteria | Enterobacteriaceae |
| Serratia sp. | -0.09684 | GCF_002607755.1 | Gammaproteobacteria | Enterobacteriaceae |
| Pseudescherichia vulneris | -0.09681 | GCF_900450975.1 | Gammaproteobacteria | Enterobacteriaceae |
| Buttiauxella sp. | -0.09678 | GCF_006376615.1 | Gammaproteobacteria | Enterobacteriaceae |
| Pantoea vagans | -0.09644 | GCF_001506165.1 | Gammaproteobacteria | Enterobacteriaceae |
| Ewingella americana | -0.09632 | GCF_000735345.1 | Gammaproteobacteria | Enterobacteriaceae |
| Pectobacterium carotovorum | -0.096 | GCF_002250215.1 | Gammaproteobacteria | Enterobacteriaceae |
| Serratia fonticola | -0.09576 | GCF_006714955.1 | Gammaproteobacteria | Enterobacteriaceae |
| Serratia sp. | -0.0957 | GCF_003668775.1 | Gammaproteobacteria | Enterobacteriaceae |
| Yersinia enterocolitica | -0.09544 | GCF_002082245.2 | Gammaproteobacteria | Enterobacteriaceae |
| Yersinia enterocolitica | -0.09536 | GCF_002083285.2 | Gammaproteobacteria | Enterobacteriaceae |
| Yersinia kristensenii | -0.09526 | GCF_002188895.1 | Gammaproteobacteria | Enterobacteriaceae |
| Morganella morganii | -0.05621 | GCF_003287815.1 | Gammaproteobacteria | Enterobacteriaceae |
| Plesiomonas sp. | -0.01279 | GCF_000800945.1 | Gammaproteobacteria | Enterobacteriaceae |
| Plesiomonas shigelloides | -0.01279 | GCF_002093895.1 | Gammaproteobacteria | Enterobacteriaceae |
| Aeromonas jandaei | -0.01248 | GCF_000708125.1 | Gammaproteobacteria | Aeromonadaceae |
| Aeromonas veronii | -0.01245 | GCF_000298015.1 | Gammaproteobacteria | Aeromonadaceae |
| Photobacterium kishitanii | -0.01235 | GCF_003025945.1 | Gammaproteobacteria | Vibrionaceae |
| Photobacterium phosphoreum | -0.01235 | GCF_003025815.1 | Gammaproteobacteria | Vibrionaceae |
| Aeromonas popoffii | -0.01224 | GCF_000820025.1 | Gammaproteobacteria | Aeromonadaceae |

**Table S3:** Top 100 over-represented annotated genes in the genomes of the taxa that receive the most negative entries on variable 1. The *NES* and *FDR-Adj. P* columns show the normalized 'Enrichment score' and FDR-adjusted (1) *P*-value from the enrichment analysis (2).

| **Gene** | **FDR-Adj. P** | **NES** |
|---|---|---|
| Major outer membrane lipoprotein Lpp 1 | -11.844 | 0.00025 |
| Phage shock protein G | -11.798 | 0.00025 |
| Primosomal replication protein N" | -11.753 | 0.00025 |
| DNA damage-inducible protein I | -11.694 | 0.00025 |
| Outer membrane porin C | -11.614 | 0.00025 |
| Chaperone protein YcdY | -11.562 | 0.00025 |
| USG-1 protein | -11.506 | 0.00025 |
| HTH-type transcriptional regulator cbl | -11.357 | 0.00025 |

| | | |
|---|---|---|
| Inner membrane protein YghB | -11.34 | 0.00025 |
| Cytochrome c-type protein NrfB | -11.321 | 0.00025 |
| Plasmid partition protein A | -11.288 | 0.00025 |
| Protein rof | -11.241 | 0.00025 |
| putative inner membrane protein Smp | -11.208 | 0.00025 |
| putative protein YbjN | -11.199 | 0.00025 |
| putative lipoprotein YbaY | -11.157 | 0.00025 |
| putative HTH-type transcriptional regulator YbdO | -11.112 | 0.00025 |
| Inner membrane protein YqjE | -11.063 | 0.00025 |
| putative protein YfeY | -11.051 | 0.00025 |
| Secretion monitor | -11.041 | 0.00025 |
| putative ECA polymerase | -11.035 | 0.00025 |
| Protein Sxy | -10.994 | 0.00025 |
| Kdo(2)-lipid A phosphoethanolamine 7"-transferase | -10.986 | 0.00025 |
| Intermembrane phospholipid transport system binding protein MlaB | -10.98 | 0.00025 |
| putative cyclic di-GMP phosphodiesterase PdeD | -10.94 | 0.00025 |
| Phosphatidylglycerophosphatase C | -10.93 | 0.00025 |
| Inner membrane protein YlaC | -10.922 | 0.00025 |
| Multiple stress resistance protein BhsA | -10.922 | 0.00025 |
| Protein YdgH | -10.922 | 0.00025 |
| Protein PhoH | -10.898 | 0.00025 |
| Outer membrane protein X | -10.882 | 0.00025 |
| Sensor protein BasS | -10.881 | 0.00025 |
| Inner membrane protein YgbE | -10.872 | 0.00025 |
| putative protein YjjI | -10.872 | 0.00025 |
| Biofilm regulator BssS | -10.87 | 0.00025 |
| DNA polymerase III subunit theta | -10.87 | 0.00025 |
| Multidrug efflux pump accessory protein AcrZ | -10.87 | 0.00025 |
| putative protein YejG | -10.87 | 0.00025 |
| putative protein YhcO | -10.87 | 0.00025 |
| Protein YhjJ | -10.864 | 0.00025 |
| Constitutive lysine decarboxylase | -10.863 | 0.00025 |
| Pirin-like protein YhaK | -10.861 | 0.00025 |
| Constitutive ornithine decarboxylase | -10.843 | 0.00025 |
| Inner membrane protein YdgK | -10.817 | 0.00025 |
| putative cyclic di-GMP phosphodiesterase PdeK | -10.815 | 0.00025 |
| Flagellar regulator flk | -10.81 | 0.00025 |
| Lipoprotein BsmA | -10.81 | 0.00025 |
| Modulator protein MzrA | -10.81 | 0.00025 |
| Inner membrane protein YgfX | -10.808 | 0.00025 |
| Protein DsrB | -10.808 | 0.00025 |
| Phosphoethanolamine transferase OpgE | -10.764 | 0.00025 |
| Transcriptional regulatory protein RcsA | -10.748 | 0.00025 |
| Osmotically-inducible lipoprotein B | -10.748 | 0.00025 |
| Cyclic di-GMP phosphodiesterase PdeH | -10.735 | 0.00025 |
| Hha toxicity modulator TomB | -10.732 | 0.00025 |
| Type II secretion system protein H | -10.728 | 0.00025 |
| putative lipoprotein YajI | -10.69 | 0.00025 |
| putative protein YebV | -10.682 | 0.00025 |
| Inner membrane protein YbjO | -10.682 | 0.00025 |
| putative lipoprotein YbjP | -10.682 | 0.00025 |
| putative protein YccJ | -10.663 | 0.00025 |
| Inner membrane protein YfeZ | -10.655 | 0.00025 |
| Flagella synthesis protein FlgN | -10.649 | 0.00025 |
| Periplasmic chaperone Spy | -10.637 | 0.00025 |
| putative ferredoxin-like protein YdhX | -10.629 | 0.00025 |
| Regulatory protein SoxS | -10.485 | 0.00025 |
| Protein TonB | -10.47 | 0.00025 |

| | | |
|---|---|---|
| Cyclic di-GMP binding protein BcsE | -10.384 | 0.00025 |
| Quorum-sensing regulator protein G | -10.381 | 0.00025 |
| Signal transduction histidine-protein kinase/phosphatase UhpB | -10.368 | 0.00025 |
| Regulator of sigma S factor FliZ | -10.328 | 0.00025 |
| Inner membrane protein YbjM | -10.323 | 0.00025 |
| Trimethylamine-N-oxide reductase | -10.318 | 0.00025 |
| Ferric iron reductase protein FhuF | -10.31 | 0.00025 |
| putative protein YaiA | -10.29 | 0.00025 |
| Negative regulator of flagellin synthesis | -10.282 | 0.00025 |
| Anti-adapter protein IraP | -10.27 | 0.00025 |
| Ferric enterobactin transport protein FepE | -10.235 | 0.00025 |
| Intracellular growth attenuator protein igaA | -10.225 | 0.00025 |
| Outer membrane porin N | -10.133 | 0.00025 |
| Flagellar protein FlhE | -10.095 | 0.00025 |
| Inner membrane protein YebE | -10.087 | 0.00025 |
| HTH-type transcriptional repressor BluR | -10.05 | 0.00025 |
| Enterobactin synthase component F | -10.035 | 0.00025 |
| Sec-independent protein translocase protein TatE | -9.992 | 0.00025 |
| Alternative ribosome-rescue factor A | -9.987 | 0.00025 |
| Primosomal protein 1 | -9.919 | 0.00025 |
| putative csgAB operon transcriptional regulatory protein | -9.909 | 0.00025 |
| putative protein YgaM | -9.887 | 0.00025 |
| Multiple antibiotic resistance protein MarA | -9.846 | 0.00025 |
| Inner membrane protein YbhQ | -9.832 | 0.00025 |
| Inner membrane protein YjiG | -9.829 | 0.00025 |
| Putative selenoprotein YdfZ | -9.827 | 0.00025 |
| HTH-type transcriptional regulator MlrA | -9.802 | 0.00025 |
| Tryptophanase | -9.8 | 0.00025 |
| Universal stress protein C | -9.798 | 0.00025 |
| Hemolysin expression-modulating protein Hha | -9.771 | 0.00025 |
| 6-phospho-beta-glucosidase BglB | -9.762 | 0.00025 |
| Ribonucleoside-diphosphate reductase 2 subunit alpha | -9.711 | 0.00025 |
| Transcriptional regulator SutA | -9.71 | 0.00025 |
| Cyclic di-GMP binding protein | -9.705 | 0.00025 |

**Table S4:** Top 100 over-represented annotated genes in the genomes of the taxa that receive the most positive entries on variable 2. The column descriptions are provided with Supplementary Table 3.

| **Gene** | **FDR-Adj. P** | **NES** |
|---|---|---|
| Outer-membrane lipoprotein LolB | 15.094 | 0.0004 |
| DNA-binding protein Fis | 15.02 | 0.0004 |
| LPS-assembly lipoprotein LptE | 15.004 | 0.0004 |
| Ammonia monooxygenase gamma subunit | 14.873 | 0.0004 |
| Thiol:disulfide interchange protein DsbA | 14.848 | 0.0004 |
| DNA polymerase III subunit delta | 14.719 | 0.0004 |
| Phosphate regulon sensor protein PhoR | 14.697 | 0.0004 |
| Pyruvate kinase II | 14.344 | 0.0004 |
| 1,6-anhydro-N-acetylmuramyl-L-alanine amidase AmpD | 14.232 | 0.0004 |
| Cell division protein FtsN | 14.223 | 0.0004 |
| Stringent starvation protein A | 14.185 | 0.0004 |
| Lipopolysaccharide export system permease protein LptF | 14.002 | 0.0004 |
| Succinate dehydrogenase hydrophobic membrane anchor subunit | 13.978 | 0.0004 |
| Recombination-associated protein RdgC | 13.944 | 0.0004 |
| Modulator of FtsH protease YccA | 13.94 | 0.0004 |
| Cell division protein ZipA | 13.785 | 0.0004 |
| Cbb3-type cytochrome c oxidase subunit CcoN1 | 13.784 | 0.0004 |
| Cytochrome c4 | 13.777 | 0.0004 |
| 2-octaprenylphenol hydroxylase | 13.656 | 0.0004 |

| | | |
|---|---|---|
| 2-octaprenyl-6-methoxyphenol hydroxylase | 13.643 | 0.0004 |
| Phosphoenolpyruvate synthase regulatory protein | 13.401 | 0.0004 |
| HTH-type transcriptional regulator CysB | 13.346 | 0.0004 |
| 2-methyl-aconitate isomerase | 13.193 | 0.0004 |
| Ferredoxin 1 | 13.12 | 0.0004 |
| Intermembrane phospholipid transport system ATP-binding protein MlaF | 13.091 | 0.0004 |
| Large ribosomal RNA subunit accumulation protein YceD | 13.051 | 0.0004 |
| Cytoskeleton protein RodZ | 13.043 | 0.0004 |
| Chorismate pyruvate-lyase | 12.981 | 0.0004 |
| Soluble lytic murein transglycosylase | 12.981 | 0.0004 |
| Intermembrane phospholipid transport system binding protein MlaD | 12.951 | 0.0004 |
| 2Fe-2S ferredoxin | 12.898 | 0.0004 |
| Cell division protein DedD | 12.872 | 0.0004 |
| Protein YcgL | 12.871 | 0.0004 |
| Uroporphyrinogen-III synthase | 12.748 | 0.0004 |
| High frequency lysogenization protein HflD | 12.712 | 0.0004 |
| putative protein YaeQ | 12.679 | 0.0004 |
| Exodeoxyribonuclease I | 12.653 | 0.0004 |
| Co-chaperone protein HscB | 12.641 | 0.0004 |
| Phosphatase NudJ | 12.606 | 0.0004 |
| Glutamate-pyruvate aminotransferase AlaA | 12.599 | 0.0004 |
| Ribonuclease T | 12.541 | 0.0004 |
| Intermembrane phospholipid transport system binding protein MlaC | 12.528 | 0.0004 |
| Colicin V production protein | 12.514 | 0.0004 |
| 5-amino-6-(5-phospho-D-ribitylamino)uracil phosphatase YigB | 12.51 | 0.0004 |
| Fimbrial protein | 12.493 | 0.0004 |
| Inner membrane transport protein YajR | 12.469 | 0.0004 |
| 50S ribosomal protein L16 3-hydroxylase | 12.462 | 0.0004 |
| Protein phosphatase CheZ | 12.411 | 0.0004 |
| FKBP-type 16 kDa peptidyl-prolyl cis-trans isomerase | 12.376 | 0.0004 |
| putative protein YibN | 12.342 | 0.0004 |
| Molybdopterin-synthase adenylyltransferase | 12.331 | 0.0004 |
| Inner membrane protein YpjD | 12.312 | 0.0004 |
| Iron-sulfur cluster assembly protein CyaY | 12.218 | 0.0004 |
| Sigma factor AlgU regulatory protein MucB | 12.217 | 0.0004 |
| Protein-glutamine gamma-glutamyltransferase | 12.205 | 0.0004 |
| Pyrimidine/purine nucleotide 5'-monophosphate nucleosidase | 12.141 | 0.0004 |
| Bifunctional (p)ppGpp synthase/hydrolase SpoT | 12.134 | 0.0004 |
| Putative glutamine amidotransferase YafJ | 12.126 | 0.0004 |
| Penicillin-binding protein 1B | 12.077 | 0.0004 |
| CDP-6-deoxy-L-threo-D-glycero-4-hexulose-3-dehydrase reductase | 12.042 | 0.0004 |
| Peptidoglycan hydrolase FlgJ | 12.011 | 0.0004 |
| Methylmalonate-semialdehyde dehydrogenase [acylating] | 11.998 | 0.0004 |
| Acid stress protein IbaG | 11.995 | 0.0004 |
| Ribosome modulation factor | 11.907 | 0.0004 |
| Cbb3-type cytochrome c oxidase subunit CcoP2 | 11.904 | 0.0004 |
| ATP synthase subunit beta 1 | 11.902 | 0.0004 |
| 3-deoxy-D-manno-octulosonic acid kinase | 11.79 | 0.0004 |
| ADP compounds hydrolase NudE | 11.777 | 0.0004 |
| HTH-type transcriptional regulator YhaJ | 11.776 | 0.0004 |
| Nucleoid-associated protein YejK | 11.714 | 0.0004 |
| Sensor protein QseC | 11.713 | 0.0004 |
| UTP pyrophosphatase | 11.706 | 0.0004 |
| ATP-dependent DNA helicase DinG | 11.685 | 0.0004 |
| Riboflavin transporter | 11.672 | 0.0004 |
| Membrane-bound lytic murein transglycosylase B | 11.611 | 0.0004 |
| putative acyltransferase YihG | 11.598 | 0.0004 |
| Protein ImuB | 11.561 | 0.0004 |

| | | |
|---|---|---|
| HTH-type transcriptional regulator HmrR | 11.528 | 0.0004 |
| DNA polymerase III subunit chi | 11.502 | 0.0004 |
| Flagellar basal-body rod protein FlgF | 11.498 | 0.0004 |
| Outer membrane protein assembly factor BamC | 11.498 | 0.0004 |
| Aerotaxis receptor | 11.478 | 0.0004 |
| Protein-glutamate methylesterase/protein-glutamine glutaminase 1 | 11.468 | 0.0004 |
| Dihydroorotase-like protein | 11.451 | 0.0004 |
| Protein YciI | 11.451 | 0.0004 |
| Cell division protein ZapD | 11.45 | 0.0004 |
| Peptidyl-prolyl cis-trans isomerase cyp18 | 11.442 | 0.0004 |
| FAD assembly factor SdhE | 11.418 | 0.0004 |
| D-erythrose-4-phosphate dehydrogenase | 11.393 | 0.0004 |
| putative DNA endonuclease SmrA | 11.37 | 0.0004 |
| Protein Smg | 11.35 | 0.0004 |
| Type II secretion system protein K | 11.347 | 0.0004 |
| Cysteine synthase A | 11.339 | 0.0004 |
| Exoribonuclease 2 | 11.332 | 0.0004 |
| Chaperone protein HscA | 11.319 | 0.0004 |
| tRNA/tmRNA (uracil-C(5))-methyltransferase | 11.319 | 0.0004 |
| Murein hydrolase activator NlpD | 11.28 | 0.0004 |
| Methyl-accepting chemotaxis protein McpP | 11.26 | 0.0004 |
| Glutathione S-transferase GST-6.0 | 11.243 | 0.0004 |
| Methyl-accepting chemotaxis protein McpH | 11.228 | 0.0004 |

**Table S5:** Top 100 over-represented annotated genes in the genomes of the taxa that receive the most positive entries on variable 3. The column descriptions are provided with Supplementary Table 3.

| **Gene** | **FDR-Adj. P** | **NES** |
|---|---|---|
| GTP pyrophosphokinase rsh | 12.285 | 0.00054 |
| putative peptidoglycan D,D-transpeptidase FtsI | 12.172 | 0.00054 |
| Chromosome-partitioning protein ParB | 12.152 | 0.00054 |
| 5-aminolevulinate synthase | 12.02 | 0.00054 |
| FtsZ-localized protein C | 11.923 | 0.00054 |
| 6,7-dimethyl-8-ribityllumazine synthase 1 | 11.812 | 0.00054 |
| FtsZ-localized protein A | 11.775 | 0.00054 |
| Aerobic cobaltochelatase subunit CobT | 11.742 | 0.00054 |
| Phyllosphere-induced regulator PhyR | 11.705 | 0.00054 |
| NADH-quinone oxidoreductase chain 1 | 11.649 | 0.00054 |
| Ubiquinone hydroxylase UbiL | 11.525 | 0.00054 |
| Cell cycle response regulator CtrA | 11.398 | 0.00054 |
| Protein phosphotransferase ChpT | 11.394 | 0.00054 |
| Heat shock protein HspQ | 11.378 | 0.00054 |
| Aerobic cobaltochelatase subunit CobS | 11.303 | 0.00054 |
| Ferredoxin-2 | 11.301 | 0.00054 |
| Dihydrolipoyl dehydrogenase 3 | 11.181 | 0.00054 |
| flagellum biosynthesis repressor protein FlbT | 11.152 | 0.00054 |
| Cytochrome c oxidase subunit 1 , bacteroid | 11.079 | 0.00054 |
| Thiol:disulfide interchange protein CycY | 11.068 | 0.00054 |
| putative protein RP812 | 10.703 | 0.00054 |
| Polyphosphate:NDP phosphotransferase 3 | 10.652 | 0.00054 |
| Serine hydroxymethyltransferase 2 | 10.633 | 0.00054 |
| Transcriptional regulatory protein ros | 10.535 | 0.00054 |
| Ferredoxin-6 | 10.444 | 0.00054 |
| Glutamate–cysteine ligase EgtA | 10.444 | 0.00054 |
| Propionyl-CoA carboxylase regulator | 10.367 | 0.00054 |
| RNA polymerase sigma-54 factor 2 | 10.354 | 0.00054 |
| Bifunctional enzyme IspD/IspF | 10.297 | 0.00054 |
| Cytochrome c1 | 10.28 | 0.00054 |

| | | |
|---|---|---|
| Cold shock protein CspA | 10.25 | 0.00054 |
| Thiol:disulfide interchange protein TlpA | 10.212 | 0.00054 |
| Nitrogen fixation regulation protein FixK | 10.079 | 0.00054 |
| Cytochrome c oxidase subunit 1-beta | 9.815 | 0.00054 |
| NADH-quinone oxidoreductase chain 5 | 9.719 | 0.00054 |
| (3S)-malyl-CoA thioesterase | 9.564 | 0.00054 |
| UDP-2,3-diacylglucosamine pyrophosphatase LpxI | 9.371 | 0.00054 |
| ATP synthase protein I | 9.331 | 0.00054 |
| Hypotaurine/taurine–pyruvate aminotransferase | 9.33 | 0.00054 |
| Blue-light-activated histidine kinase | 9.329 | 0.00054 |
| HTH-type transcriptional regulator RamB | 9.327 | 0.00054 |
| Porin | 9.325 | 0.00054 |
| Penicillin-insensitive murein endopeptidase | 9.315 | 0.00054 |
| Glycine betaine methyltransferase | 9.294 | 0.00054 |
| L-arabinose 1-dehydrogenase (NAD(P)(+)) | 9.267 | 0.00054 |
| Putative metal-sulfur cluster biosynthesis proteins YuaD | 9.264 | 0.00054 |
| ATP synthase subunit b’ | 9.25 | 0.00054 |
| N-acetylmuramoyl-L-alanine amidase AmiD | 9.212 | 0.00054 |
| Periplasmic alpha-galactoside-binding protein | 9.195 | 0.00054 |
| 10 kDa chaperonin 1 | 9.186 | 0.00054 |
| Urease subunit gamma 1 | 9.152 | 0.00054 |
| (2S)-methylsuccinyl-CoA dehydrogenase | 9.097 | 0.00054 |
| Urease subunit alpha 1 | 9.066 | 0.00054 |
| Hemolysin C | 9.047 | 0.00054 |
| Urease accessory protein UreE 1 | 8.894 | 0.00054 |
| Lysine/ornithine decarboxylase | 8.874 | 0.00054 |
| Dicamba O-demethylase 1, ferredoxin reductase component | 8.873 | 0.00054 |
| Serine–glyoxylate aminotransferase | 8.855 | 0.00054 |
| Precorrin-3B C(17)-methyltransferase | 8.849 | 0.00054 |
| Nicotinate phosphoribosyltransferase | 8.819 | 0.00054 |
| Glycogen synthase 1 | 8.812 | 0.00054 |
| 3-hydroxybenzoate 6-hydroxylase 1 | 8.768 | 0.00054 |
| nicotinate-nucleotide adenylyltransferase | 8.741 | 0.00054 |
| Molybdenum cofactor insertion chaperone PaoD | 8.735 | 0.00054 |
| Arginine–pyruvate transaminase AruH | 8.718 | 0.00054 |
| D-hydantoinase/dihydropyrimidinase | 8.695 | 0.00054 |
| putative 3-hydroxyisobutyrate dehydrogenase | 8.592 | 0.00054 |
| Response regulator receiver protein CpdR | 8.56 | 0.00054 |
| 60 kDa chaperonin 5 | 8.547 | 0.00054 |
| Lysophospholipase L2 | 8.545 | 0.00054 |
| Type I secretion system ATP-binding protein PrsD | 8.544 | 0.00054 |
| Phosphatidylcholine synthase | 8.531 | 0.00054 |
| Carbonic anhydrase 1 | 8.507 | 0.00054 |
| Bifunctional coenzyme PQQ synthesis protein C/D | 8.503 | 0.00054 |
| NAD-dependent dihydropyrimidine dehydrogenase subunit PreT | 8.474 | 0.00054 |
| Anti-sigma-F factor NrsF | 8.456 | 0.00054 |
| FAD-dependent catabolic D-arginine dehydrogenase DauA | 8.437 | 0.00054 |
| Nopaline-binding periplasmic protein | 8.432 | 0.00054 |
| Mesaconyl-CoA hydratase | 8.431 | 0.00054 |
| putative riboflavin import permease protein RfuD | 8.419 | 0.00054 |
| Sulfite dehydrogenase subunit C | 8.391 | 0.00054 |
| Crotonyl-CoA carboxylase/reductase | 8.381 | 0.00054 |
| Precorrin-2 C(20)-methyltransferase | 8.38 | 0.00054 |
| Acyl carrier protein AcpXL | 8.376 | 0.00054 |
| Polysialic acid transport ATP-binding protein KpsT | 8.372 | 0.00054 |
| Alkane 1-monooxygenase 2 | 8.352 | 0.00054 |
| HTH-type transcriptional regulator RafR | 8.331 | 0.00054 |
| S-formylglutathione hydrolase | 8.326 | 0.00054 |

| | | |
|---|---|---|
| Ethylmalonyl-CoA mutase | 8.282 | 0.00054 |
| Outer membrane protein | 8.273 | 0.00054 |
| Cytochrome c oxidase subunit 4 | 8.252 | 0.00054 |
| Cytochrome c-556 | 8.237 | 0.00054 |
| Alpha-D-ribose 1-methylphosphonate 5-triphosphate synthase subunit PhnG | 8.236 | 0.00054 |
| Alpha-D-ribose 1-methylphosphonate 5-triphosphate diphosphatase | 8.223 | 0.00054 |
| Sulfite dehydrogenase subunit A | 8.218 | 0.00054 |
| Glutathione-specific gamma-glutamylcyclotransferase | 8.17 | 0.00054 |
| (S)-ureidoglycine aminohydrolase | 8.165 | 0.00054 |
| Hydrogenobyrinate a,c-diamide synthase | 8.101 | 0.00054 |
| Alanine racemase, biosynthetic | 8.068 | 0.00054 |
| Alpha-D-ribose 1-methylphosphonate 5-triphosphate synthase subunit PhnH | 8.045 | 0.00054 |

**Table S6:** Top 100 over-represented annotated genes in the genomes of the taxa that receive the most negative entries on variable 4. The column descriptions are provided with Supplementary Table 3.

| Gene | FDR-Adj. P | NES |
|---|---|---|
| Cytochrome f | -6.889 | 0.00031 |
| Photosystem II manganese-stabilizing polypeptide | -6.876 | 0.00031 |
| Protein Thf1 | -6.874 | 0.00031 |
| Photosystem II CP47 reaction center protein | -6.859 | 0.00031 |
| NAD(P)H-quinone oxidoreductase subunit N | -6.857 | 0.00031 |
| Photosystem I assembly protein Ycf4 | -6.857 | 0.00031 |
| Photosystem I reaction center subunit III | -6.857 | 0.00031 |
| Phycocyanobilin:ferredoxin oxidoreductase | -6.85 | 0.00031 |
| Photosystem II reaction center Psb28 protein | -6.836 | 0.00031 |
| Photosystem II lipoprotein Psb27 | -6.835 | 0.00031 |
| NAD(P)H-quinone oxidoreductase subunit O | -6.833 | 0.00031 |
| Photosystem I P700 chlorophyll a apoprotein A1 | -6.833 | 0.00031 |
| Photosystem II reaction center protein K | -6.829 | 0.00031 |
| Cytochrome b559 subunit alpha | -6.826 | 0.00031 |
| Photosystem II CP43 reaction center protein | -6.826 | 0.00031 |
| Photosystem I reaction center subunit IV | -6.824 | 0.00031 |
| 30S ribosomal protein S21 A | -6.812 | 0.00031 |
| Pentapeptide repeat protein Rfr32 | -6.807 | 0.00031 |
| Photosystem II reaction center protein H | -6.804 | 0.00031 |
| Ycf54-like protein | -6.803 | 0.00031 |
| Ferredoxin-thioredoxin reductase, catalytic chain | -6.802 | 0.00031 |
| NAD(P)H-quinone oxidoreductase subunit M | -6.795 | 0.00031 |
| Long-chain acyl-[acyl-carrier-protein] reductase | -6.795 | 0.00031 |
| Bifunctional pantoate ligase/cytidylate kinase | -6.792 | 0.00031 |
| RNA polymerase sigma factor SigA2 | -6.78 | 0.00031 |
| Photosystem I reaction center subunit II | -6.779 | 0.00031 |
| Photosystem I reaction center subunit XI | -6.779 | 0.00031 |
| Phycobiliprotein beta chain | -6.773 | 0.00031 |
| Phycobilisome 7.8 kDa linker polypeptide, allophycocyanin-associated, core | -6.773 | 0.00031 |
| Photosystem I reaction center subunit XII | -6.769 | 0.00031 |
| Aldehyde decarbonylase | -6.752 | 0.00031 |
| Photosystem II reaction center protein Z | -6.75 | 0.00031 |
| Ferredoxin-thioredoxin reductase, variable chain | -6.739 | 0.00031 |
| Photosystem II 12 kDa extrinsic protein | -6.73 | 0.00031 |
| Photosystem II protein Y | -6.723 | 0.00031 |
| Protein PsbN | -6.723 | 0.00031 |
| Proton extrusion protein PcxA | -6.719 | 0.00031 |
| Cytochrome b6-f complex subunit 7 | -6.698 | 0.00031 |
| Phycocyanobilin lyase subunit alpha | -6.692 | 0.00031 |
| Orange carotenoid-binding protein | -6.684 | 0.00031 |
| Cytochrome b559 subunit beta | -6.665 | 0.00031 |

| | | |
|---|---|---|
| Photosystem I iron-sulfur center | -6.638 | 0.00031 |
| Vitamin K epoxide reductase | -6.628 | 0.00031 |
| Photosystem I reaction center subunit IX | -6.626 | 0.00031 |
| ATP-dependent zinc metalloprotease FtsH 2 | -6.612 | 0.00031 |
| Putative diflavin flavoprotein A 3 | -6.577 | 0.00031 |
| putative glutaredoxin | -6.556 | 0.00031 |
| Monoglucosyldiacylglycerol epimerase | -6.546 | 0.00031 |
| Photosystem II protein D1 2 | -6.546 | 0.00031 |
| NAD(P)H-quinone oxidoreductase subunit L | -6.543 | 0.00031 |
| Lipoyl synthase 2 | -6.517 | 0.00031 |
| Phycocyanobilin lyase CpcT | -6.495 | 0.00031 |
| 2-methyl-6-phytyl-1,4-hydroquinone methyltransferase | -6.492 | 0.00031 |
| Photosystem II reaction center X protein | -6.491 | 0.00031 |
| Photosystem II D2 protein | -6.47 | 0.00031 |
| Transcription regulator LexA | -6.463 | 0.00031 |
| Ycf53-like protein | -6.463 | 0.00031 |
| D-fructose 1,6-bisphosphatase class 2/sedoheptulose 1,7-bisphosphatase | -6.447 | 0.00031 |
| putative arabinosyltransferase C | -6.445 | 0.00031 |
| Phycobiliprotein ApcE | -6.435 | 0.00031 |
| Photosystem II reaction center protein T | -6.434 | 0.00031 |
| NAD(P)H-quinone oxidoreductase subunit K 1 | -6.424 | 0.00031 |
| Sensor protein SphS | -6.419 | 0.00031 |
| Allophycocyanin beta chain | -6.412 | 0.00031 |
| Photosystem II reaction center protein M | -6.396 | 0.00031 |
| Photosystem II reaction center protein Ycf12 | -6.39 | 0.00031 |
| Circadian clock protein KaiA | -6.388 | 0.00031 |
| Photosystem I reaction center subunit VIII | -6.371 | 0.00031 |
| High-affinity Na(+)/H(+) antiporter NhaS3 | -6.34 | 0.00031 |
| putative 30S ribosomal protein PSRP-3 | -6.308 | 0.00031 |
| Isoaspartyl peptidase/L-asparaginase | -6.297 | 0.00031 |
| Phytol kinase | -6.261 | 0.00031 |
| Regulatory protein CysR | -6.248 | 0.00031 |
| Photosystem II reaction center protein I | -6.243 | 0.00031 |
| Serine/threonine-protein kinase B | -6.237 | 0.00031 |
| Ferredoxin-dependent glutamate synthase 2 | -6.229 | 0.00031 |
| Phycobilisome rod-core linker polypeptide CpcG | -6.226 | 0.00031 |
| Galactan 5-O-arabinofuranosyltransferase | -6.217 | 0.00031 |
| NAD(P)H-quinone oxidoreductase subunit J | -6.213 | 0.00031 |
| Putative serine protease HhoA | -6.174 | 0.00031 |
| Phycocyanobilin lyase subunit CpcS | -6.151 | 0.00031 |
| Photosystem II reaction center protein J | -6.149 | 0.00031 |
| Putative acetyl-coenzyme A carboxylase carboxyl transferase subunit beta | -6.134 | 0.00031 |
| Phycocyanobilin lyase subunit beta | -6.125 | 0.00031 |
| NAD(P)H-quinone oxidoreductase subunit I | -6.123 | 0.00031 |
| Phosphoribulokinase | -6.118 | 0.00031 |
| Putative isochorismate synthase MenF | -6.103 | 0.00031 |
| Allophycocyanin subunit alpha-B | -6.097 | 0.00031 |
| Putative cytochrome P450 120 | -6.075 | 0.00031 |
| Putative diflavin flavoprotein A 5 | -6.071 | 0.00031 |
| Bicarbonate-binding protein CmpA | -6.071 | 0.00031 |
| C-phycocyanin beta chain | -6.068 | 0.00031 |
| Chromophore lyase CpcS/CpeS | -6.065 | 0.00031 |
| 4-hydroxybenzoate solanesyltransferase | -6.034 | 0.00031 |
| Hydrolase | -6.024 | 0.00031 |
| L,D-transpeptidase 2 | -6.004 | 0.00031 |
| putative ferredoxin/ferredoxin–NADP reductase | -5.991 | 0.00031 |
| Photosystem II reaction center protein L | -5.972 | 0.00031 |
| Alpha-(1-¿3)-arabinofuranosyltransferase | -5.963 | 0.00031 |

| | | |
|---|---|---|
| Serine/threonine-protein kinase F | -5.943 | 0.00031 |

**Table S7:** Top 100 over-represented annotated genes in the genomes of the taxa that receive the most negative entries on variable 14. The column descriptions are provided with Supplementary Table 3.

| Gene | FDR-Adj. P | NES |
|---|---|---|
| Thymidylate synthase 1 | -5.522 | 0.00055 |
| DNA-binding protein Bv3F | -5.446 | 0.00055 |
| mupirocin-resistant isoleucine–tRNA ligase MupA | -5.294 | 0.00055 |
| Cobalt-precorrin-7 C(5)-methyltransferase | -5.114 | 0.00055 |
| Cobalt-zinc-cadmium resistance protein CzcI | -5.103 | 0.00055 |
| IS200/IS605 family transposase ISCth10 | -4.897 | 0.00055 |
| Na(+)-translocating ferredoxin:NAD(+) oxidoreductase complex subunit C | -4.843 | 0.00055 |
| Anaerobic sulfite reductase subunit C | -4.814 | 0.00055 |
| (R)-2-hydroxyglutaryl-CoA dehydratase activating ATPase | -4.812 | 0.00055 |
| Salicylate 5-hydroxylase, large oxygenase component | -4.778 | 0.00055 |
| Tyrosine aminotransferase | -4.765 | 0.00055 |
| putative protein YgcP | -4.703 | 0.00055 |
| Alkaline phosphatase PhoK | -4.685 | 0.00055 |
| Salicylate 5-hydroxylase, small oxygenase component | -4.684 | 0.00055 |
| Propanediol utilization protein PduU | -4.64 | 0.00055 |
| Putative superoxide reductase | -4.596 | 0.00055 |
| Sortase B | -4.552 | 0.00055 |
| Propanediol utilization protein PduV | -4.424 | 0.00055 |
| Elongation factor G, mitochondrial | -4.343 | 0.00055 |
| Germination protease | -4.315 | 0.00055 |
| Stage IV sporulation protein A | -4.315 | 0.00055 |
| Stage V sporulation protein AD | -4.315 | 0.00055 |
| putative N-glycosylase/DNA lyase | -4.309 | 0.00055 |
| (R)-phenyllactyl-CoA dehydratase alpha subunit | -4.297 | 0.00055 |
| Nickel-cobalt-cadmium resistance protein NccX | -4.295 | 0.00055 |
| (R)-2-hydroxyglutaryl-CoA dehydratase, subunit beta | -4.293 | 0.00055 |
| Putative transport protein YbjL | -4.292 | 0.00055 |
| RNA polymerase sigma-G factor | -4.286 | 0.00055 |
| Translocation-enhancing protein TepA | -4.271 | 0.00055 |
| Stage V sporulation protein T | -4.261 | 0.00055 |
| Light-activated DNA-binding protein EL222 | -4.254 | 0.00055 |
| RNA polymerase sigma-28 factor | -4.245 | 0.00055 |
| putative anti-sigma-F factor NrsF | -4.242 | 0.00055 |
| Oxalate-binding protein | -4.225 | 0.00055 |
| IS256 family transposase ISCth4 | -4.222 | 0.00055 |
| Propanediol utilization protein PduB | -4.202 | 0.00055 |
| Stage III sporulation protein D | -4.195 | 0.00055 |
| Spore protein YabP | -4.194 | 0.00055 |
| 3,4-dehydroadipyl-CoA semialdehyde dehydrogenase | -4.18 | 0.00055 |
| Nickel and cobalt resistance protein CnrR | -4.118 | 0.00055 |
| Neopullulanase 1 | -4.117 | 0.00055 |
| Small, acid-soluble spore protein C2 | -4.111 | 0.00055 |
| Mini-ribonuclease 3-like protein | -4.103 | 0.00055 |
| L-threonine kinase | -4.09 | 0.00055 |
| IS3 family transposase ISStma17 | -4.039 | 0.00055 |
| Outer membrane protein 40 | -4.031 | 0.00055 |
| Methionine-rich peptide X | -3.993 | 0.00055 |
| IS5 family transposase ISBmu20 | -3.97 | 0.00055 |
| Reverse rubrerythrin-1 | -3.967 | 0.00055 |
| Histidine racemase | -3.951 | 0.00055 |
| Spore germination protein B1 | -3.947 | 0.00055 |
| 2-pyrone-4,6-dicarboxylate hydrolase | -3.936 | 0.00055 |

| | | |
|---|---|---|
| Glycine/sarcosine/betaine reductase complex component C subunit alpha | -3.931 | 0.00055 |
| Tryptophanase 1 | -3.919 | 0.00055 |
| Nickel and cobalt resistance protein CnrC | -3.918 | 0.00055 |
| putative sporulation protein YlmC | -3.91 | 0.00055 |
| mupirocin-resistant isoleucine–tRNA ligase MupB | -3.907 | 0.00055 |
| putative deoxyuridine 5'-triphosphate nucleotidohydrolase YncF | -3.89 | 0.00055 |
| IS110 family transposase ISCaa14 | -3.886 | 0.00055 |
| SpoIVB peptidase | -3.885 | 0.00055 |
| Glycine reductase complex component B subunit gamma | -3.874 | 0.00055 |
| Propionate catabolism operon regulatory protein | -3.87 | 0.00055 |
| RNA polymerase sigma-35 factor | -3.865 | 0.00055 |
| Propanediol dehydratase medium subunit | -3.852 | 0.00055 |
| Chloroacetanilide N-alkylformylase, ferredoxin reductase component | -3.835 | 0.00055 |
| Ribulose bisphosphate carboxylase large chain, chromosomal | -3.835 | 0.00055 |
| IS66 family transposase ISBcen14 | -3.824 | 0.00055 |
| putative tryptophan transport protein | -3.82 | 0.00055 |
| IS66 family transposase ISBcen19 | -3.809 | 0.00055 |
| Peptidoglycan-N-acetylmuramic acid deacetylase PdaA | -3.808 | 0.00055 |
| Light-harvesting protein B-870 beta chain | -3.798 | 0.00055 |
| PEP-dependent dihydroxyacetone kinase 2, phosphoryl donor subunit DhaM | -3.796 | 0.00055 |
| Glycine reductase complex component B subunits alpha and beta | -3.793 | 0.00055 |
| IS1182 family transposase ISCpe5 | -3.79 | 0.00055 |
| Accessory gene regulator protein B | -3.788 | 0.00055 |
| CRISPR-associated endoribonuclease Cas6 | -3.787 | 0.00055 |
| Phosphoglycolate phosphatase, plasmid | -3.786 | 0.00055 |
| Glycerol dehydratase large subunit | -3.781 | 0.00055 |
| Glycine/sarcosine/betaine reductase complex component C subunit beta | -3.773 | 0.00055 |
| Diol dehydratase-reactivating factor alpha subunit | -3.759 | 0.00055 |
| IS3 family transposase ISElsp1 | -3.754 | 0.00055 |
| Stage II sporulation protein E | -3.754 | 0.00055 |
| IS110 family transposase ISCaa7 | -3.754 | 0.00055 |
| Propanediol dehydratase small subunit | -3.748 | 0.00055 |
| Reverse rubrerythrin-2 | -3.747 | 0.00055 |
| Cytochrome c-type protein SHP | -3.744 | 0.00055 |
| Antigen TpF1 | -3.731 | 0.00055 |
| Serine/threonine-protein kinase CtkA | -3.731 | 0.00055 |
| Diadenosine hexaphosphate hydrolase | -3.73 | 0.00055 |
| Glycine/sarcosine/betaine reductase complex component A | -3.724 | 0.00055 |
| Outer membrane protein 41 | -3.721 | 0.00055 |
| Stage III sporulation protein AE | -3.703 | 0.00055 |
| N-acetylmuramoyl-L-alanine amidase | -3.7 | 0.00055 |
| Catechol 1,2-dioxygenase 2 | -3.696 | 0.00055 |
| Glycine/sarcosine/betaine reductase complex component A1 | -3.678 | 0.00055 |
| D-proline reductase proprotein PrdA | -3.654 | 0.00055 |
| Catechol 1,2-dioxygenase 1 | -3.633 | 0.00055 |
| Phthalate 4,5-dioxygenase oxygenase reductase subunit | -3.63 | 0.00055 |
| Iron hydrogenase 1 | -3.623 | 0.00055 |
| Metal-staphylopine import system ATP-binding protein CntD | -3.618 | 0.00055 |

**Table S8:** Top 100 over-represented annotated genes in the genomes of the taxa that receive the most negative entries on variable 38. The column descriptions are provided with Supplementary Table 3.

| **Gene** | **FDR-Adj. P** | **NES** |
|---|---|---|
| Bifunctional protein MdtA | -5.039 | 0.00064 |
| Flagellar assembly protein FliX | -4.91 | 0.00064 |
| Presqualene diphosphate synthase | -4.54 | 0.00064 |
| mupirocin-resistant isoleucine–tRNA ligase MupA | -4.537 | 0.00064 |
| Formyltransferase/hydrolase complex subunit D | -4.476 | 0.00064 |

| | | |
|---|---|---|
| Formyltransferase/hydrolase complex Fhc subunit A | -4.455 | 0.00064 |
| Methenyltetrahydromethanopterin cyclohydrolase | -4.406 | 0.00064 |
| 3',5'-cyclic-nucleotide phosphodiesterase | -4.329 | 0.00064 |
| Formyltransferase/hydrolase complex Fhc subunit C | -4.301 | 0.00064 |
| Methylmalonyl-CoA mutase small subunit | -4.277 | 0.00064 |
| Bifunctional dihydropteroate synthase/dihydropteroate reductase | -4.27 | 0.00064 |
| Plasminogen-binding protein PgbB | -4.237 | 0.00064 |
| Oxygen-independent coproporphyrinogen-III oxidase-like protein HemZ | -4.205 | 0.00064 |
| GTP cyclohydrolase 1 type 2 | -4.195 | 0.00064 |
| 5,6,7,8-tetrahydromethanopterin hydro-lyase | -4.19 | 0.00064 |
| Methanol dehydrogenase [cytochrome c] subunit 2 | -4.181 | 0.00064 |
| Sensor protein DivL | -4.173 | 0.00064 |
| Hydroxycarboxylate dehydrogenase B | -4.148 | 0.00064 |
| Sortase B | -4.142 | 0.00064 |
| 2-amino-5-chloromuconate deaminase | -4.127 | 0.00064 |
| L-hydantoinase | -4.024 | 0.00064 |
| Cytochrome c-L | -4.022 | 0.00064 |
| Flagellar FliL protein | -4.012 | 0.00064 |
| Putative ATP-dependent DNA helicase YjcD | -3.986 | 0.00064 |
| Beta-methylmalyl-CoA dehydratase | -3.968 | 0.00064 |
| Cytochrome c-553 | -3.959 | 0.00064 |
| (2R)-sulfolactate sulfo-lyase subunit alpha | -3.955 | 0.00064 |
| Malyl-CoA/beta-methylmalyl-CoA/citramalyl-CoA lyase | -3.953 | 0.00064 |
| Oxalate:formate antiporter | -3.934 | 0.00064 |
| Inducible ornithine decarboxylase | -3.921 | 0.00064 |
| Dihydromethanopterin reductase | -3.891 | 0.00064 |
| 10 kDa chaperonin 2 | -3.884 | 0.00064 |
| Na(+)-translocating ferredoxin:NAD(+) oxidoreductase complex subunit G | -3.882 | 0.00064 |
| Lipoprotein NlpI | -3.873 | 0.00064 |
| Bifunctional DNA-directed RNA polymerase subunit beta-beta' | -3.83 | 0.00064 |
| Inner membrane protein YabI | -3.823 | 0.00064 |
| Cbb3-type cytochrome c oxidase subunit FixP | -3.797 | 0.00064 |
| Bifunctional coenzyme PQQ synthesis protein C/D | -3.774 | 0.00064 |
| Opacity-associated protein OapA | -3.769 | 0.00064 |
| Surface-adhesin protein E | -3.769 | 0.00064 |
| Accessory gene regulator protein B | -3.752 | 0.00064 |
| Blue-light absorbing proteorhodopsin | -3.709 | 0.00064 |
| Beta-(1–¿2)glucan export ATP-binding/permease protein NdvA | -3.708 | 0.00064 |
| Chaperone protein YcdY | -3.702 | 0.00064 |
| Ribulose bisphosphate carboxylase large chain 2 | -3.684 | 0.00064 |
| Outer membrane protein P5 | -3.68 | 0.00064 |
| Chloramphenicol resistance protein CraA | -3.654 | 0.00064 |
| Na(+)-translocating ferredoxin:NAD(+) oxidoreductase complex subunit C | -3.644 | 0.00064 |
| Neopullulanase 1 | -3.64 | 0.00064 |
| DNA transformation protein TfoX | -3.632 | 0.00064 |
| Beta-carotene 15,15'-dioxygenase | -3.631 | 0.00064 |
| Redox-sensing transcriptional repressor Rex 1 | -3.631 | 0.00064 |
| Metallopeptidase AprA | -3.629 | 0.00064 |
| Rubrerythrin-1 | -3.628 | 0.00064 |
| Hydrogenase/urease maturation factor HypB | -3.615 | 0.00064 |
| Small, acid-soluble spore protein C2 | -3.598 | 0.00064 |
| Ribulose bisphosphate carboxylase small chain 2 | -3.595 | 0.00064 |
| Translocation-enhancing protein TepA | -3.591 | 0.00064 |
| Gamma-glutamyl-L-1-hydroxyisopropylamide hydrolase | -3.587 | 0.00064 |
| putative protein YgcP | -3.566 | 0.00064 |
| RNA polymerase sigma-28 factor | -3.526 | 0.00064 |
| PTS system N-acetylglucosamine-specific EIIB component | -3.52 | 0.00064 |
| Na(+)-translocating ferredoxin:NAD(+) oxidoreductase complex subunit D | -3.513 | 0.00064 |

| | | |
|---|---|---|
| Germination protease | -3.511 | 0.00064 |
| Stage IV sporulation protein A | -3.511 | 0.00064 |
| Squalene–hopene cyclase | -3.508 | 0.00064 |
| putative cobalt-factor III C(17)-methyltransferase | -3.491 | 0.00064 |
| USG-1 protein | -3.485 | 0.00064 |
| Valine dehydrogenase | -3.482 | 0.00064 |
| Molybdenum storage protein subunit alpha | -3.479 | 0.00064 |
| Stage V sporulation protein AD | -3.476 | 0.00064 |
| RNA polymerase sigma-G factor | -3.474 | 0.00064 |
| 5-(methylthio)ribulose-1-phosphate aldolase | -3.473 | 0.00064 |
| Translational regulator CsrA2 | -3.469 | 0.00064 |
| Spore protein YabP | -3.459 | 0.00064 |
| Stage V sporulation protein T | -3.446 | 0.00064 |
| Oxalyl-CoA decarboxylase | -3.444 | 0.00064 |
| Quinone-reactive Ni/Fe-hydrogenase large chain | -3.442 | 0.00064 |
| Translational regulator CsrA1 | -3.44 | 0.00064 |
| L-proline trans-4-hydroxylase | -3.437 | 0.00064 |
| Undecaprenyl-diphosphooligosaccharide–protein glycotransferase | -3.428 | 0.00064 |
| D(-)-tartrate dehydratase | -3.423 | 0.00064 |
| Resuscitation-promoting factor Rpf | -3.42 | 0.00064 |
| Nucleoid-associated protein Lsr2 | -3.405 | 0.00064 |
| Elongation factor G, mitochondrial | -3.399 | 0.00064 |
| Potassium/sodium uptake protein NtpJ | -3.398 | 0.00064 |
| Malate synthase | -3.396 | 0.00064 |
| Formyltransferase/hydrolase complex Fhc subunit B | -3.394 | 0.00064 |
| Stage III sporulation protein D | -3.381 | 0.00064 |
| Putative septation protein SpoVG | -3.369 | 0.00064 |
| Hemolysin C | -3.365 | 0.00064 |
| Tyrosine recombinase XerH | -3.355 | 0.00064 |
| NAD(P)-dependent methylenetetrahydromethanopterin dehydrogenase | -3.355 | 0.00064 |
| Oxalate decarboxylase OxdD | -3.353 | 0.00064 |
| Cobalt-dependent inorganic pyrophosphatase | -3.352 | 0.00064 |
| 60 kDa chaperonin 3 | -3.351 | 0.00064 |
| Protein PhoH | -3.35 | 0.00064 |
| Glutathione amide-dependent peroxidase | -3.342 | 0.00064 |
| putative quinol monooxygenase YgiN | -3.326 | 0.00064 |
| DNA-binding protein HB1 | -3.326 | 0.00064 |

**Table S9:** Top 100 over-represented annotated genes in the genomes of the taxa that receive the most positive entries on variable 43. The column descriptions are provided with Supplementary Table 3.

| **Gene** | **FDR-Adj. P** | **NES** |
|---|---|---|
| Sirohydrochlorin cobaltochelatase CbiKP | 5.05 | 0.00064 |
| Bifunctional protein MdtA | 4.982 | 0.00064 |
| Cytochrome c-L | 4.843 | 0.00064 |
| Methanol dehydrogenase [cytochrome c] subunit 2 | 4.798 | 0.00064 |
| Carbon monoxide dehydrogenase 1 | 4.71 | 0.00064 |
| NAD(+)–dinitrogen-reductase ADP-D-ribosyltransferase | 4.631 | 0.00064 |
| Carbon monoxide dehydrogenase/acetyl-CoA synthase subunit alpha | 4.487 | 0.00064 |
| Corrinoid/iron-sulfur protein large subunit | 4.47 | 0.00064 |
| Hydrogenase-2 large chain | 4.455 | 0.00064 |
| Molybdenum storage protein subunit beta | 4.406 | 0.00064 |
| Sulfite reductase, dissimilatory-type subunit gamma | 4.308 | 0.00064 |
| Acetolactate synthase isozyme 1 small subunit | 4.265 | 0.00064 |
| Protein DsvD | 4.254 | 0.00064 |
| mupirocin-resistant isoleucine–tRNA ligase MupA | 4.214 | 0.00064 |
| Hopanoid C-3 methylase | 4.162 | 0.00064 |
| Menaquinone reductase, iron-sulfur cluster-binding subunit | 4.161 | 0.00064 |

| | | |
|---|---|---|
| Metal-binding protein SmbP | 4.145 | 0.00064 |
| Menaquinone reductase, molybdopterin-binding-like subunit | 4.065 | 0.00064 |
| Reverse rubrerythrin-1 | 4.054 | 0.00064 |
| Rubredoxin 3 | 4.026 | 0.00064 |
| Hydrogenase-2 small chain | 3.991 | 0.00064 |
| Ribulose bisphosphate carboxylase small chain 2 | 3.969 | 0.00064 |
| Periplasmic [NiFe] hydrogenase large subunit | 3.965 | 0.00064 |
| Ribulose bisphosphate carboxylase large chain 2 | 3.918 | 0.00064 |
| Menaquinone reductase, multiheme cytochrome c subunit | 3.913 | 0.00064 |
| (R)-2-hydroxyisocaproyl-CoA dehydratase beta subunit | 3.895 | 0.00064 |
| Formyltransferase/hydrolase complex Fhc subunit B | 3.822 | 0.00064 |
| Toluene-4-monooxygenase system, ferredoxin component | 3.812 | 0.00064 |
| Hydroxylamine oxidoreductase | 3.79 | 0.00064 |
| Menaquinone reductase, integral membrane subunit | 3.777 | 0.00064 |
| Accessory gene regulator protein B | 3.775 | 0.00064 |
| Resuscitation-promoting factor Rpf | 3.76 | 0.00064 |
| Split-Soret cytochrome c | 3.752 | 0.00064 |
| Carbon monoxide dehydrogenase 2 | 3.747 | 0.00064 |
| Dihydromethanopterin reductase | 3.726 | 0.00064 |
| Protein FeSII | 3.712 | 0.00064 |
| PTS system N-acetylglucosamine-specific EIIB component | 3.705 | 0.00064 |
| Nitrogen fixation regulatory protein | 3.626 | 0.00064 |
| Na(+)-translocating ferredoxin:NAD(+) oxidoreductase complex subunit C | 3.608 | 0.00064 |
| Alpha-amylase 1 | 3.584 | 0.00064 |
| Elongation factor G, mitochondrial | 3.576 | 0.00064 |
| Neopullulanase 1 | 3.574 | 0.00064 |
| Hydrogenase-4 component G | 3.525 | 0.00064 |
| Ribulose bisphosphate carboxylase | 3.525 | 0.00064 |
| D-xylonate dehydratase YagF | 3.51 | 0.00064 |
| Sporulation-specific cell division protein SsgB | 3.506 | 0.00064 |
| EtfAB:quinone oxidoreductase | 3.505 | 0.00064 |
| Lipoprotein NlpI | 3.494 | 0.00064 |
| CRISPR-associated endonuclease Cas6 | 3.461 | 0.00064 |
| Cytochrome c-type protein ImcH | 3.421 | 0.00064 |
| Opacity-associated protein OapA | 3.417 | 0.00064 |
| Surface-adhesin protein E | 3.417 | 0.00064 |
| Fused nickel transport protein NikMN | 3.416 | 0.00064 |
| Molybdenum storage protein subunit alpha | 3.414 | 0.00064 |
| Corrinoid/iron-sulfur protein small subunit | 3.399 | 0.00064 |
| Small, acid-soluble spore protein C2 | 3.399 | 0.00064 |
| Cytochrome c" | 3.384 | 0.00064 |
| IS1182 family transposase ISRssp12 | 3.375 | 0.00064 |
| Mannosylglucosyl-3-phosphoglycerate synthase | 3.347 | 0.00064 |
| Flagellar FliL protein | 3.344 | 0.00064 |
| Valine dehydrogenase | 3.333 | 0.00064 |
| Sensor protein CseC | 3.319 | 0.00064 |
| Tyrosine-protein kinase CpsD | 3.312 | 0.00064 |
| Rubredoxin-oxygen oxidoreductase | 3.306 | 0.00064 |
| putative nitrate/nitrite transporter NarK2 | 3.298 | 0.00064 |
| 5-hydroxybenzimidazole synthase BzaA | 3.297 | 0.00064 |
| Enoyl-[acyl-carrier-protein] reductase [NADPH] FabI | 3.295 | 0.00064 |
| Putative sulfur carrier protein YeeD | 3.292 | 0.00064 |
| Translocation-enhancing protein TepA | 3.28 | 0.00064 |
| IS66 family transposase ISSwo2 | 3.258 | 0.00064 |
| Citrate (Re)-synthase | 3.257 | 0.00064 |
| putative protein YgcP | 3.257 | 0.00064 |
| PTS system N-acetylglucosamine-specific EIIC component | 3.256 | 0.00064 |
| Putative superoxide reductase | 3.256 | 0.00064 |

| | | |
|---|---|---|
| Cyanuric acid amidohydrolase | 3.247 | 0.00064 |
| IS91 family transposase ISCARN110 | 3.24 | 0.00064 |
| IS5 family transposase ISPso2 | 3.234 | 0.00064 |
| IS1595 family transposase ISMpo2 | 3.232 | 0.00064 |
| 60 kDa chaperonin 3 | 3.229 | 0.00064 |
| Particulate methane monooxygenase beta subunit | 3.206 | 0.00064 |
| Oxalate oxidoreductase subunit beta | 3.205 | 0.00064 |
| 10 kDa chaperonin 2 | 3.2 | 0.00064 |
| Sucrose synthase | 3.199 | 0.00064 |
| putative sporulation protein YlmC | 3.187 | 0.00064 |
| IS66 family transposase ISDpr4 | 3.186 | 0.00064 |
| DNA transformation protein TfoX | 3.185 | 0.00064 |
| (R)-2-hydroxyisocaproyl-CoA dehydratase alpha subunit | 3.17 | 0.00064 |
| Outer membrane protein P5 | 3.159 | 0.00064 |
| putative secretion system apparatus ATP synthase SsaN | 3.15 | 0.00064 |
| IS1182 family transposase ISClbu1 | 3.15 | 0.00064 |
| 2-amino-5-chloromuconate deaminase | 3.147 | 0.00064 |
| Type A flavoprotein fprA | 3.145 | 0.00064 |
| Benzylsuccinate synthase activating enzyme | 3.144 | 0.00064 |
| (R)-phenyllactate dehydratase activator | 3.142 | 0.00064 |
| Particulate methane monooxygenase alpha subunit | 3.133 | 0.00064 |
| (R)-phenyllactyl-CoA dehydratase alpha subunit | 3.126 | 0.00064 |
| Barbiturase 1 | 3.122 | 0.00064 |
| CRISPR system Cascade subunit CasE | 3.112 | 0.00064 |
| NADH-dependent phenylglyoxylate dehydrogenase subunit gamma | 3.106 | 0.00064 |
| ECF RNA polymerase sigma factor ShbA | 3.103 | 0.00064 |

### Supplementary Figures

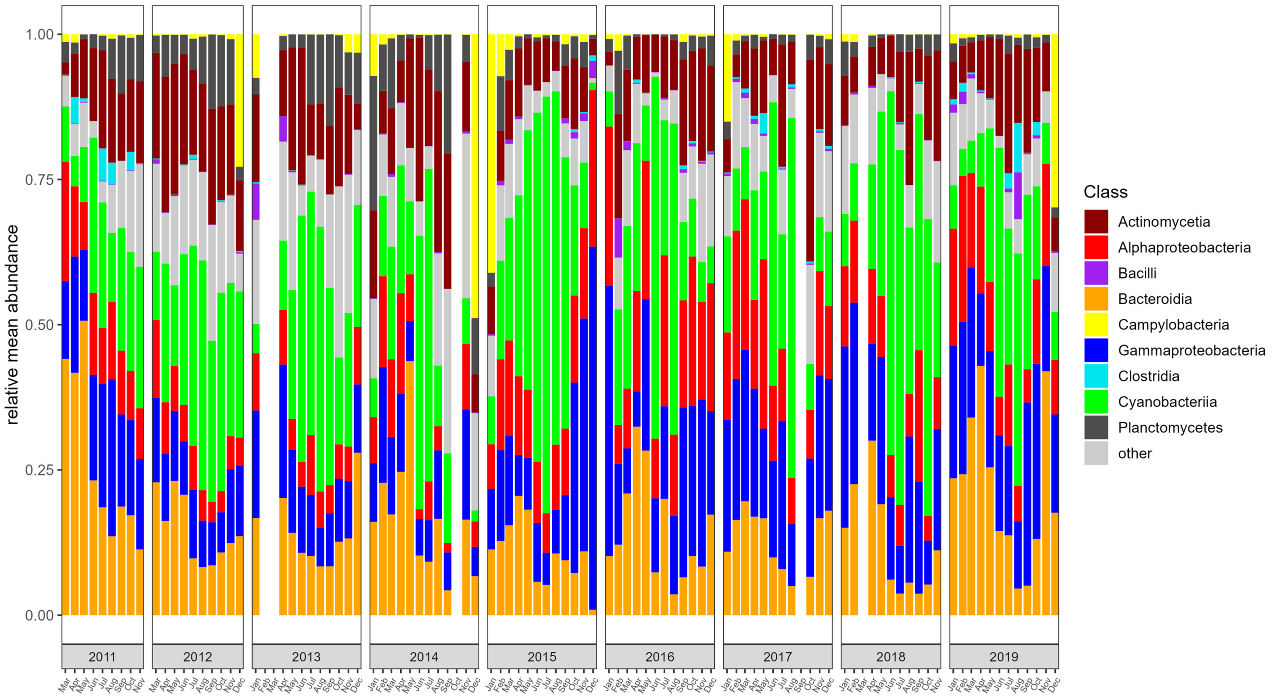


**Figure S1:** Relative mean abundances of classes that map to the genomes obtained from amplicon sequencing data over the whole sampling period. Taxonomic classes are color-coded.

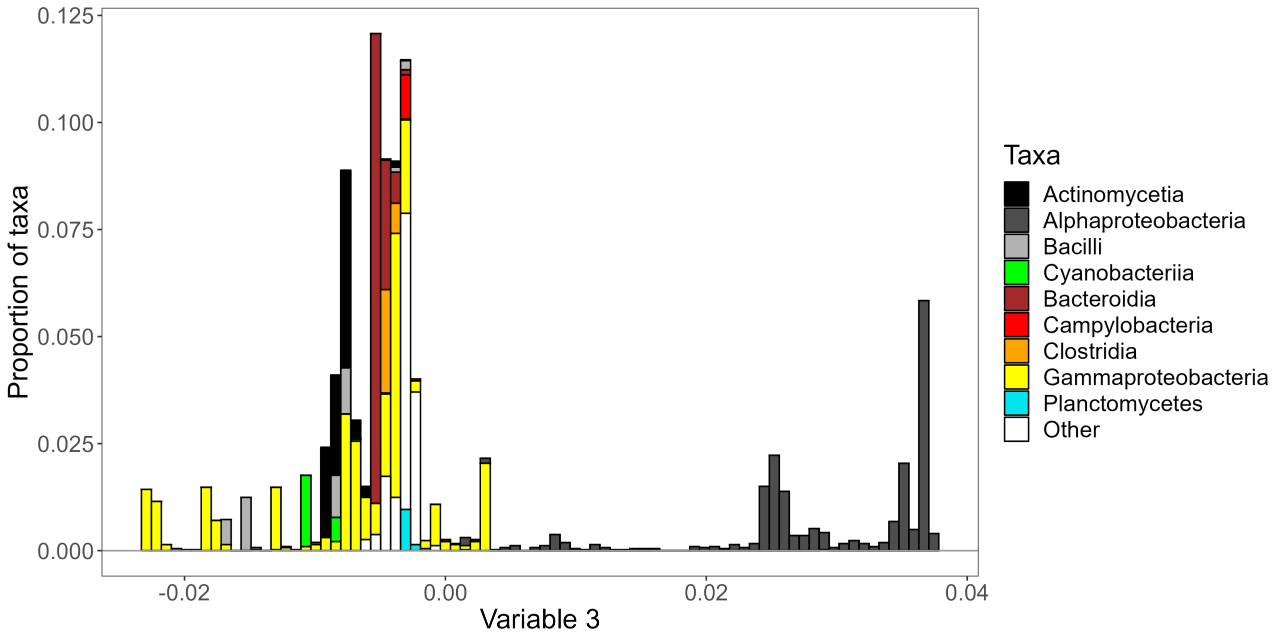


**Figure S2:** The ordering of taxa defined by variable 3 entries, from negative to positive (left to right). The taxonomic compositions corresponding to variable entries are shown for each of 80 equally spaced bins.

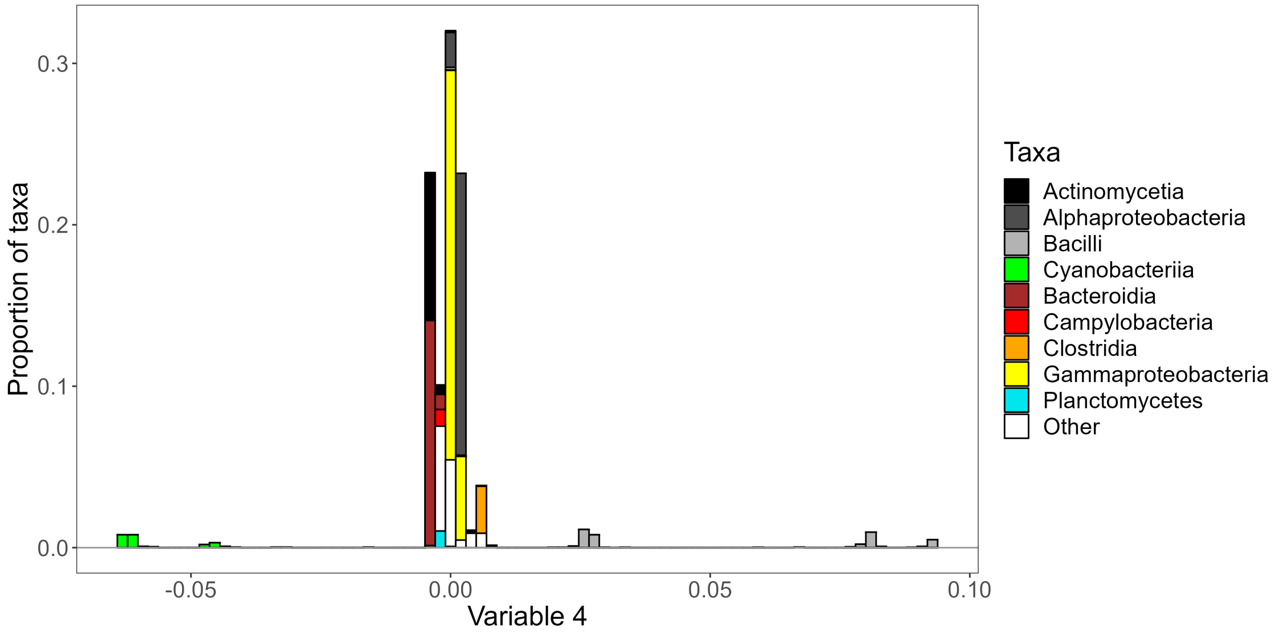


**Figure S3:** The ordering of taxa defined by variable 4 entries, from negative to positive (left to right). The taxonomic compositions corresponding to variable entries are shown for each of 80 equally spaced bins.

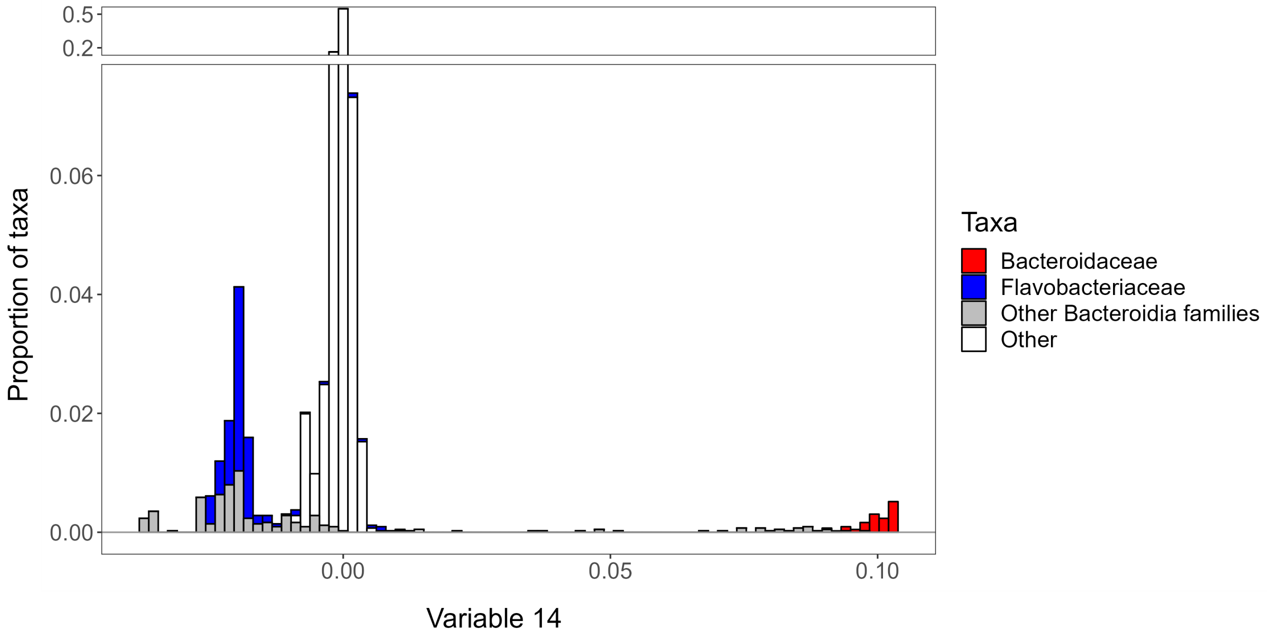


**Figure S4:** The ordering of taxa defined by variable 14 entries, from negative to positive (left to right). The taxonomic compositions corresponding to variable entries are shown for each of 80 equally spaced bins. Families belonging to the phylum Bacteroidota are color-coded.

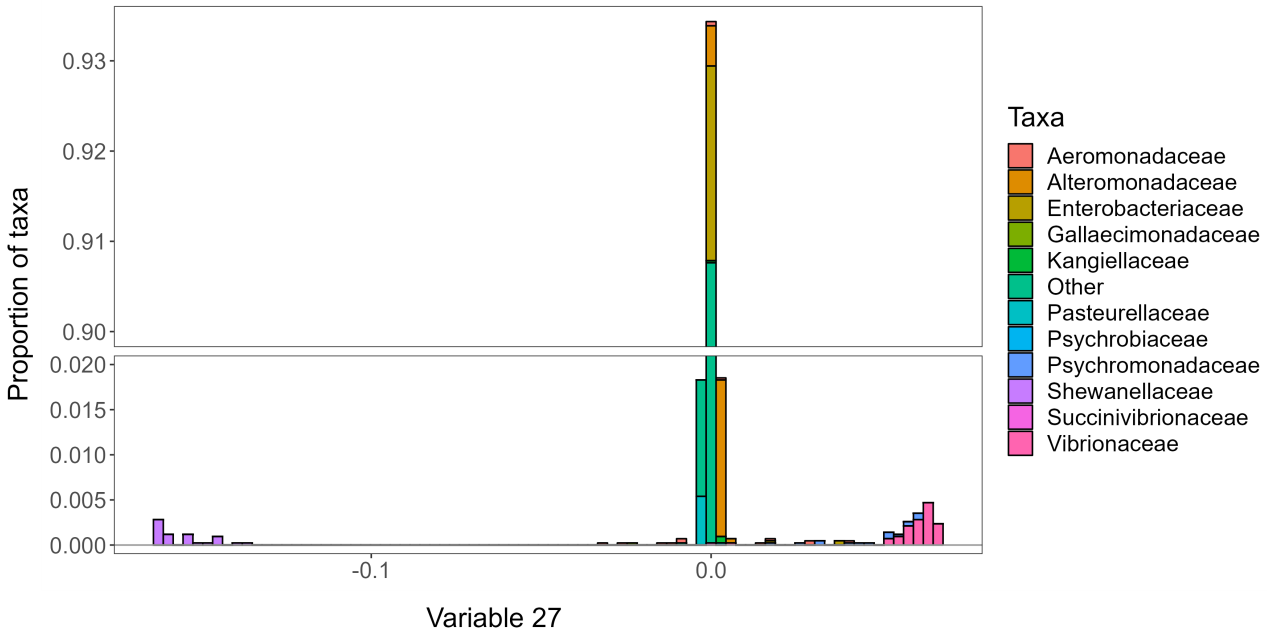


**Figure S5:** The ordering of taxa defined by variable 27 entries, from negative to positive (left to right). The taxonomic compositions corresponding to variable entries are shown for each of 80 equally spaced bins. Families belonging to the Enterobacterales are color-coded.

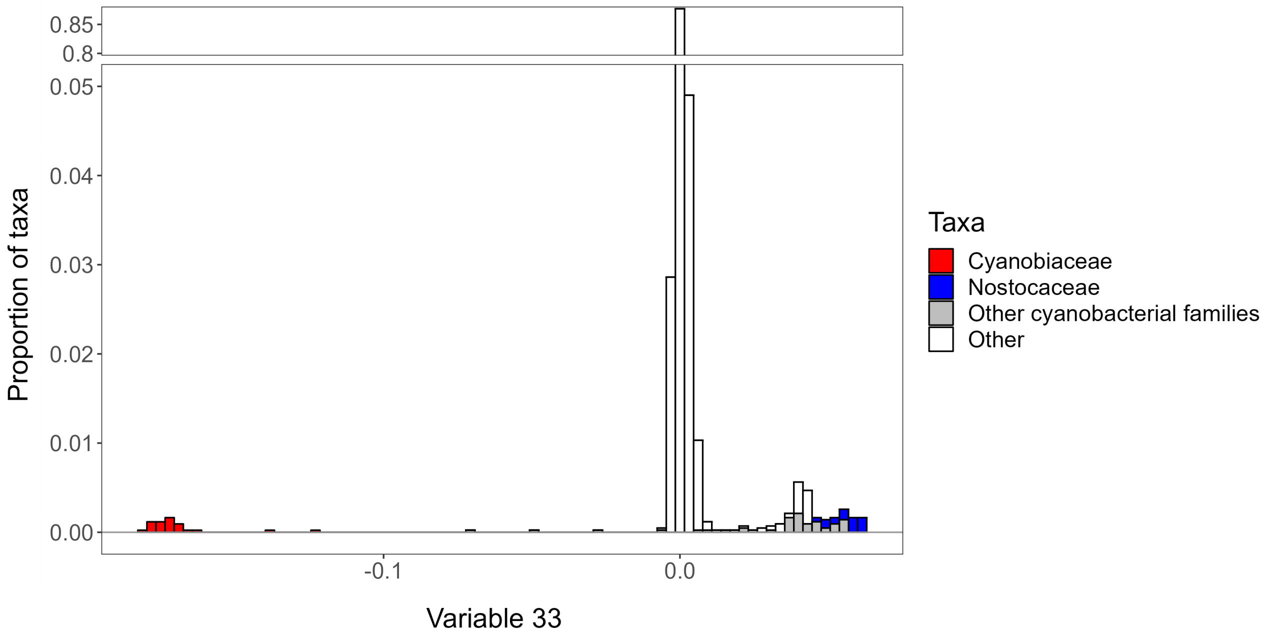


**Figure S6:** The ordering of taxa defined by variable 33 entries, from negative to positive (left to right). The taxonomic compositions corresponding to variable entries are shown for each of 80 equally spaced bins. Families belonging to the class of Cyanobacteriia are color-coded.

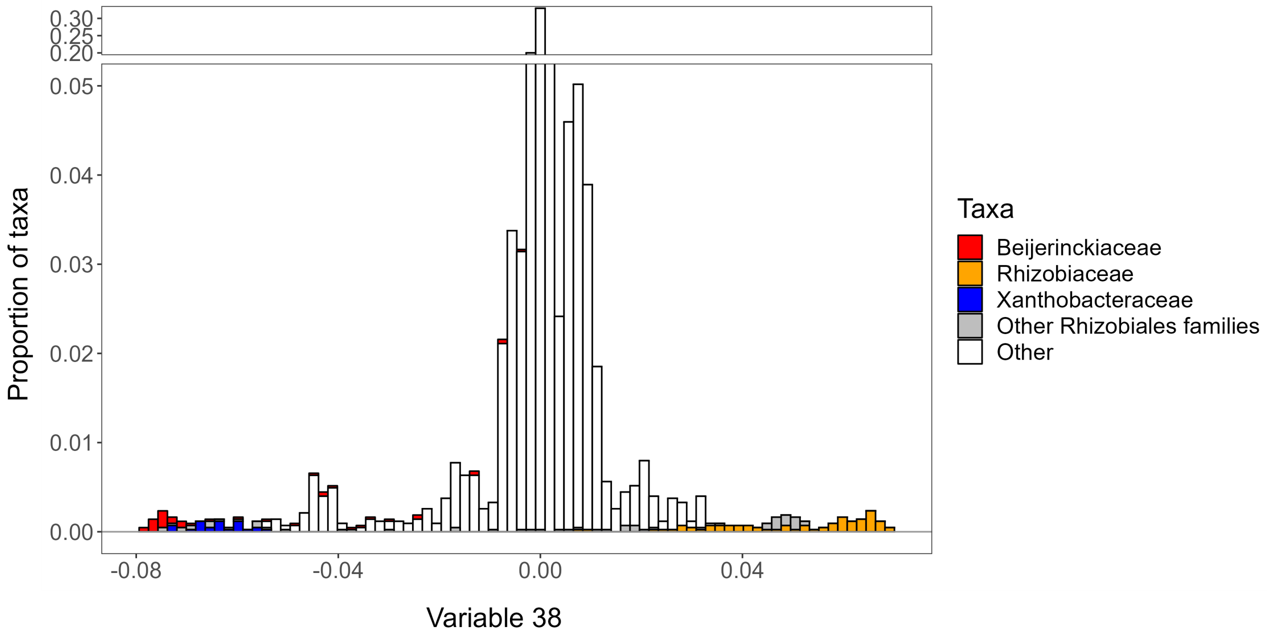


**Figure S7:** The ordering of taxa defined by variable 38 entries, from negative to positive (left to right). The taxonomic compositions corresponding to variable entries are shown for each of 80 equally spaced bins. Families belonging to the Order Rhizobiales are color-coded.

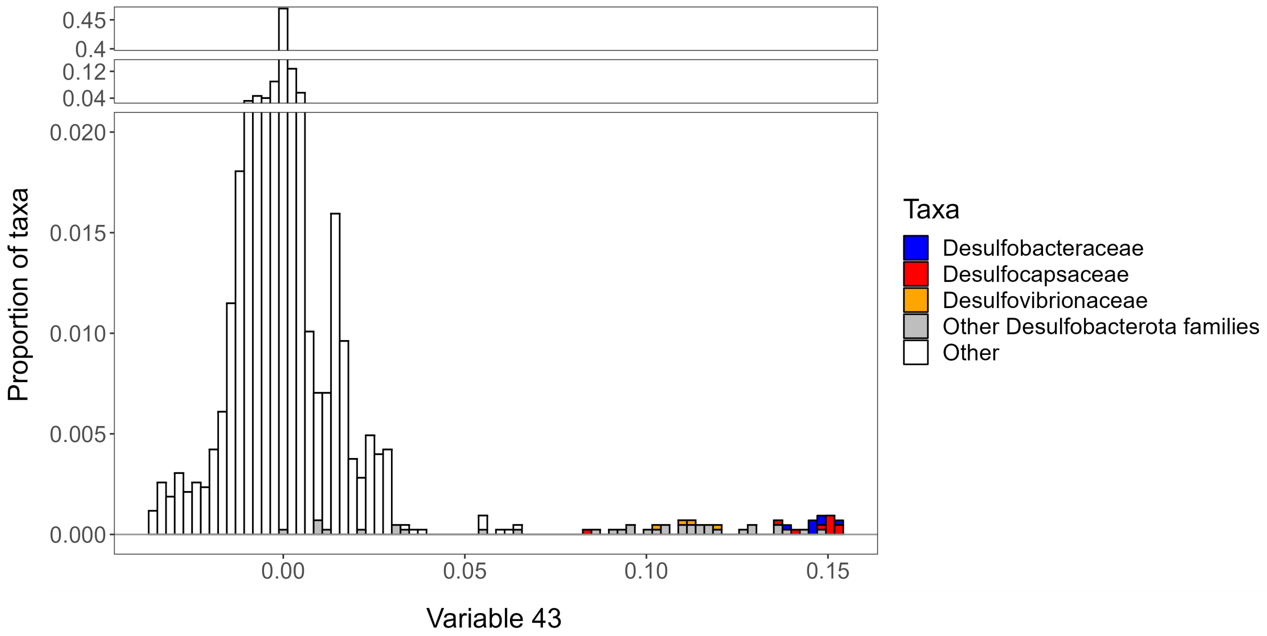


**Figure S8:** The ordering of taxa defined by variable 43 entries, from negative to positive (left to right). The taxonomic compositions corresponding to variable entries are shown for each of 80 equally spaced bins. Families belonging to the phylum Desulfobacterota are color-coded.

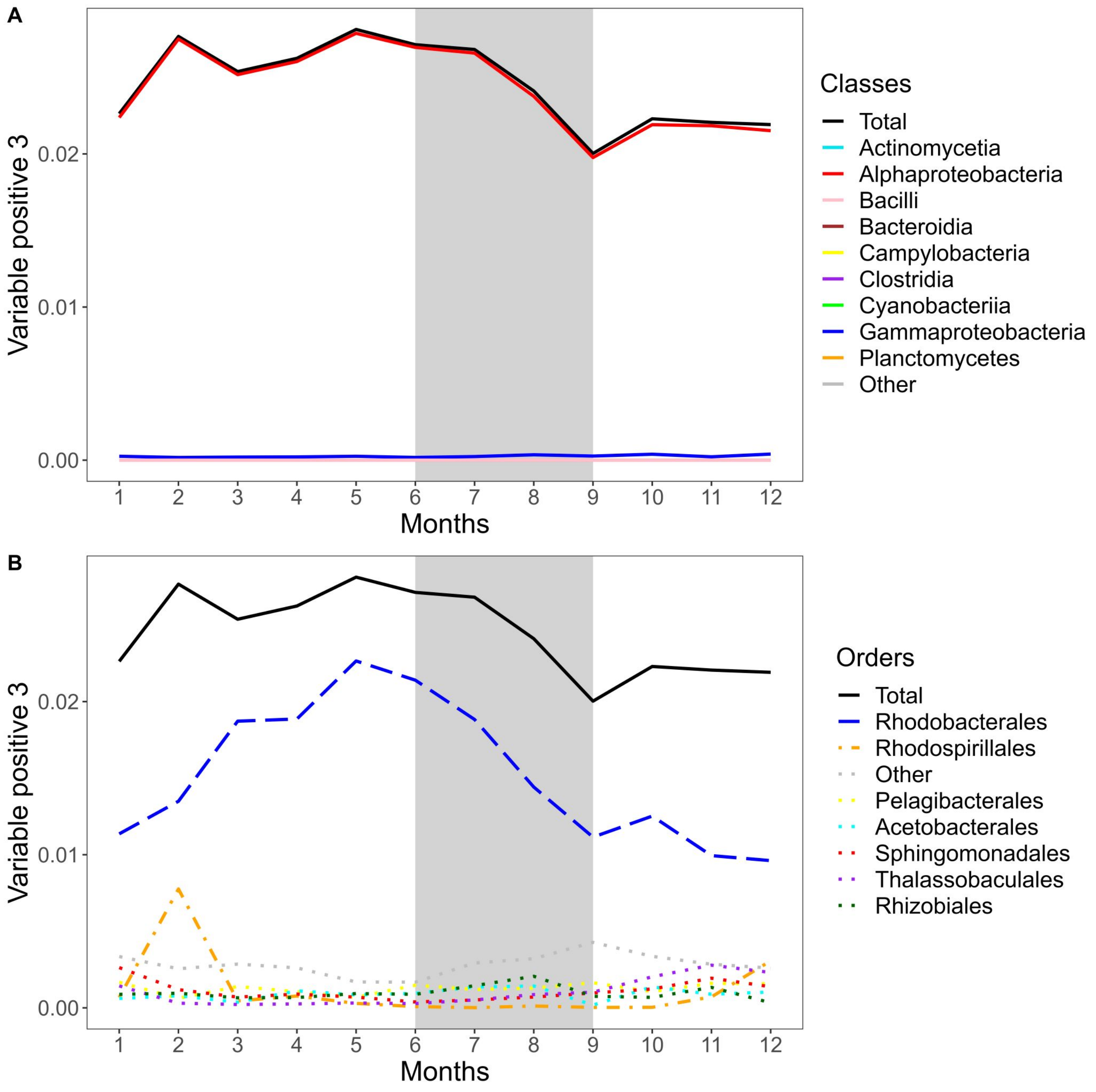


**Figure S9:** Abundance-weighted mean values of inferred ability of utilizing a variety of carbon sources over the yearly cycle. Summer months are indicated by a gray background. Taxonomic class (A) and taxonomic orders (B) are color-coded.

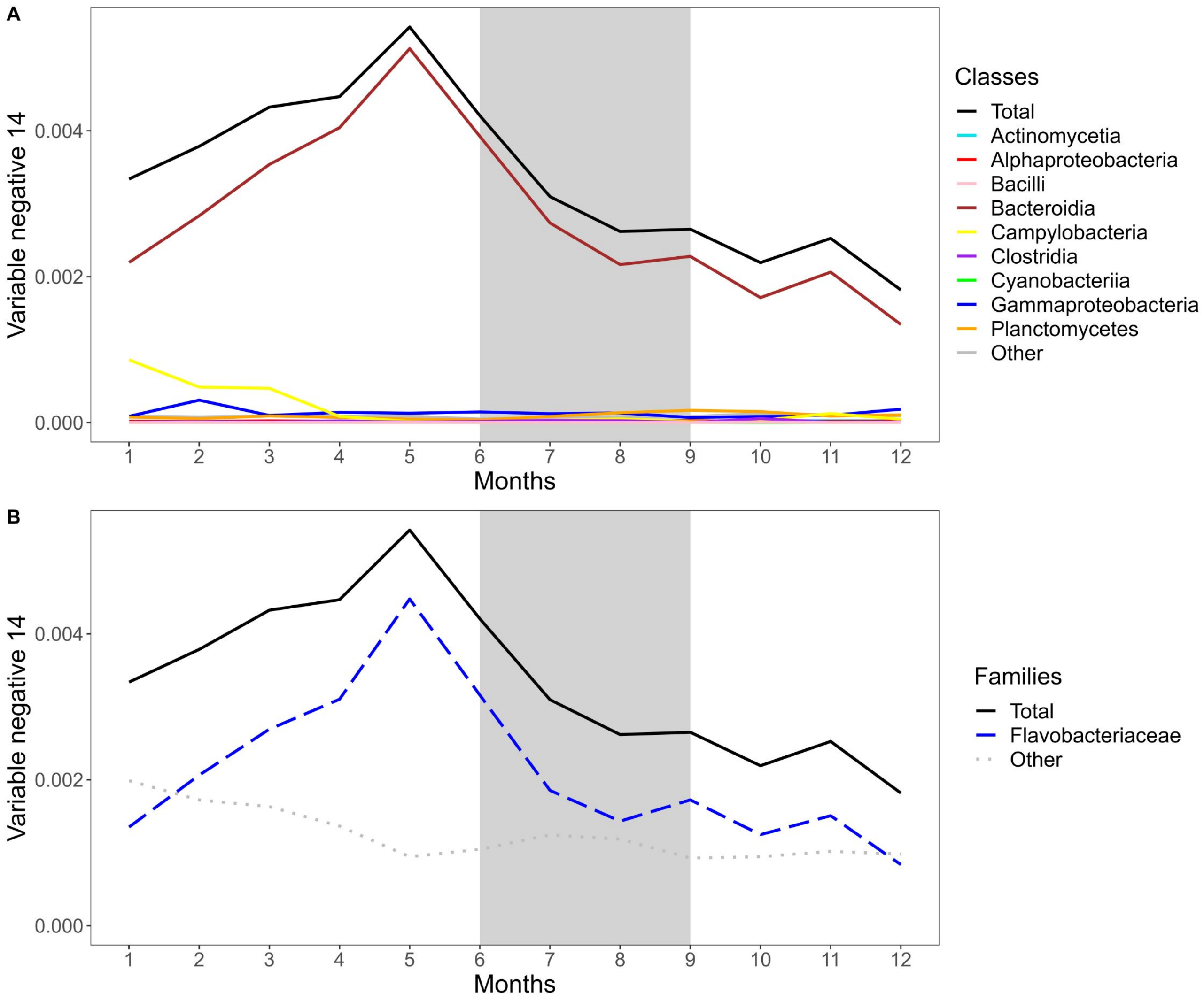


**Figure S10:** Abundance-weighted mean values of inferred ability of degrading complex polysaccharides over the yearly cycle. Summer months are indicated by a gray background. Taxonomic class (A) and taxonomic families (B) are color-coded.

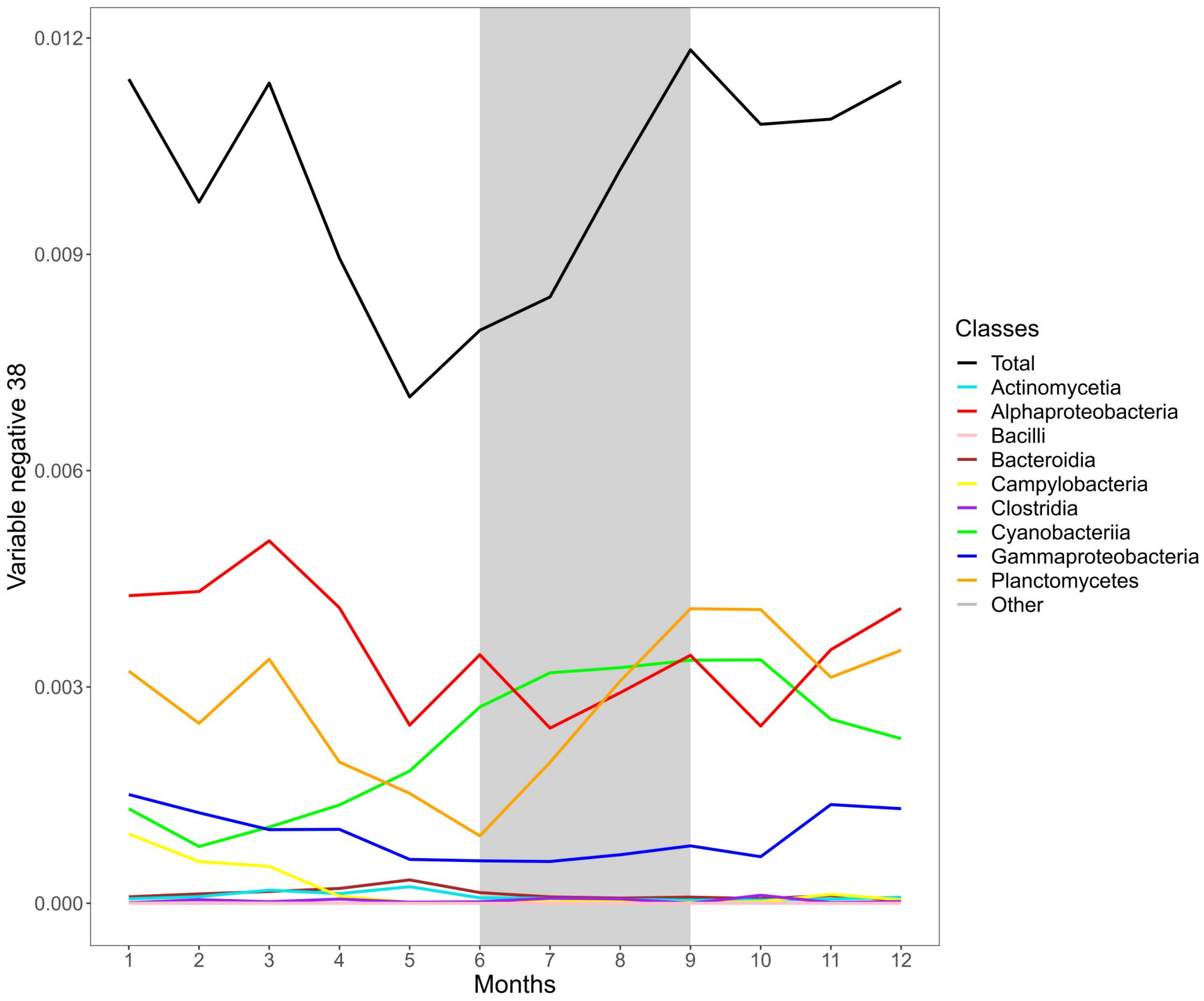


**Figure S11:** Abundance-weighted mean values of inferred ability of oxidizing methyl groups and C1 compounds over the yearly cycle. Summer months are indicated by a gray background. Taxonomic class is color-coded.

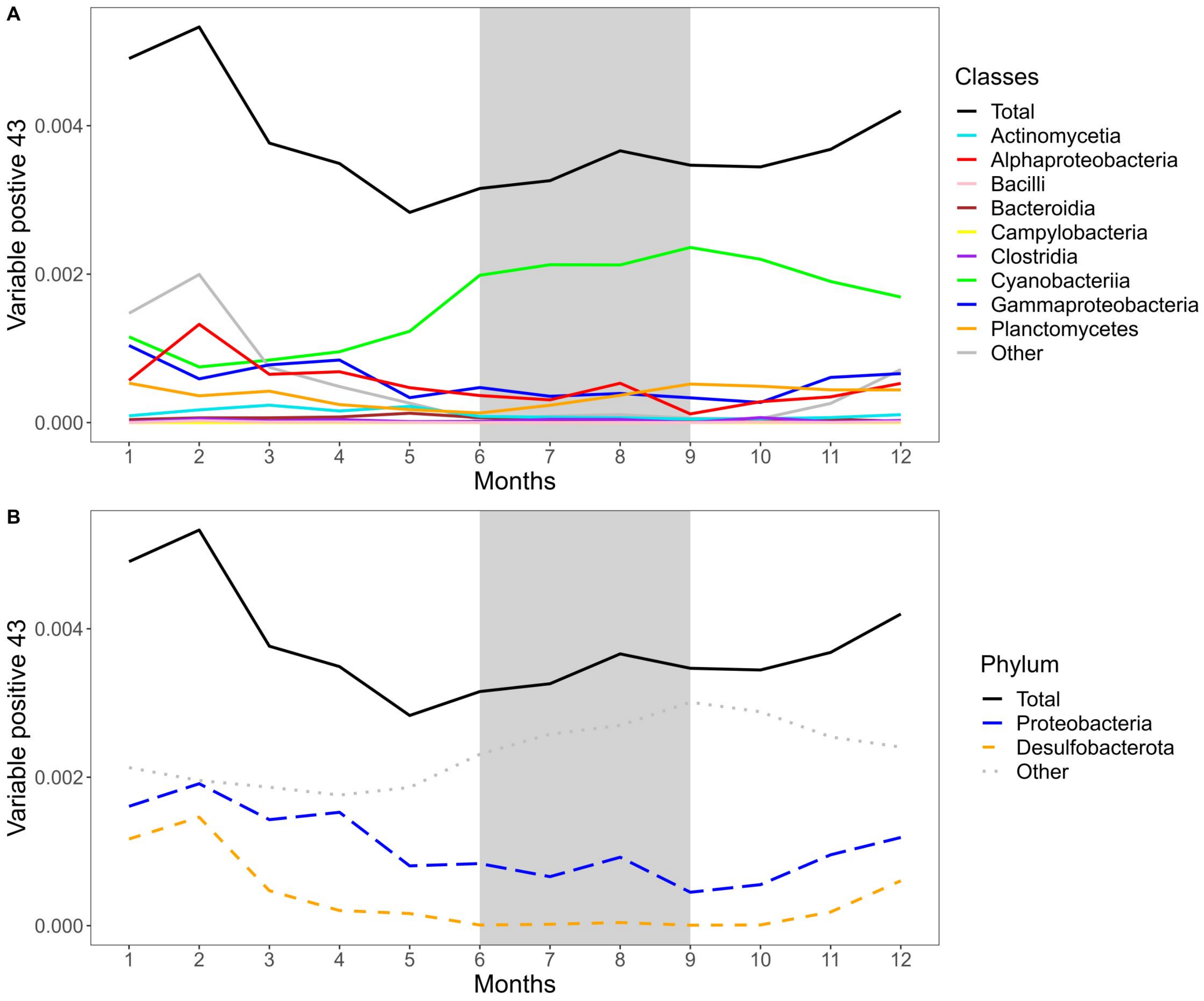


**Figure S12:** Abundance-weighted mean values of trait dominated by non-spore forming sulfate reducers over the yearly cycle. Summer months are indicated by a gray background. Taxonomic class is color-coded.

### References

[1] Yoav Benjamini and Yosef Hochberg. Controlling the false discovery rate: a practical and powerful approach to multiple testing. *Journal of the Royal statistical society: series B (Methodological)*, 57(1):289–300, 1995.

[2] Aravind Subramanian, Pablo Tamayo, Vamsi K Mootha, Sayan Mukherjee, Benjamin L Ebert, Michael A Gillette, Amanda Paulovich, Scott L Pomeroy, Todd R Golub, Eric S Lander, et al. Gene set enrichment analysis: a knowledge-based approach for interpreting genome-wide expression profiles. *Proceedings of the National Academy of Sciences*, 102(43):15545–15550, 2005.
